# Cellular proteostasis decline in human senescence

**DOI:** 10.1101/860775

**Authors:** Niv Sabath, Flonia Levy-Adam, Amal Younis, Kinneret Rozales, Anatoly Meller, Shani Hadar, Sharon Soueid-Baumgarten, Reut Shalgi

**Affiliations:** Department of Biochemistry, Rappaport Faculty of Medicine, Technion–Israel Institute of Technology, Haifa 31096, Israel

**Keywords:** Protein homeostasis, chaperones, UPR, HSF1, Heat shock response, Senescence

## Abstract

Proteostasis collapse, the diminished ability to maintain protein homeostasis, has been established as a hallmark of nematode aging. However, whether proteostasis collapse occurs in humans has remained unclear. Here we demonstrate that proteostasis decline is intrinsic to human senescence. Using transcriptome-wide characterization of gene expression, splicing and translation, we found a significant deterioration in the transcriptional activation of the heat shock response in stressed senescent cells. Furthermore, phosphorylated HSF1 nuclear localization and distribution were impaired in senescence. Interestingly, alternative splicing regulation was also dampened. Surprisingly, we found a decoupling between different Unfolded Protein Response (UPR) branches in stressed senescent cells. While young cells initiated UPR-related translational and transcriptional regulatory responses, senescent cells showed enhanced translational regulation and ER stress sensing, however they were unable to trigger UPR-related transcriptional responses. This was accompanied by diminished ATF6 nuclear localization in stressed senescent cells. Finally, we revealed a deterioration of proteasome function in senescence following heat stress, which did not recover upon return to normal temperature. Together, our data unraveled a deterioration in the ability to mount dynamic stress transcriptional programs upon human senescence with broad implications on proteostasis control, and connected proteostasis decline to human aging.

**Significance:** Protein homeostasis (proteostasis), the balance between protein synthesis, folding, and degradation, is thought to deteriorate with age, and the prevalence of protein misfolding diseases, e.g. Alzheimer’s, Parkinson’s etc., with human aging is increased. However, while in worms this phenomenon has been well established, in humans it remained unclear. Here we show that proteostasis is declined in human senescence, i.e. cellular aging. We found that while stress sensing is enhanced in senescent cells, and their response at the level of protein synthesis is intact, they fail to properly activate multiple programs required for stress adaptation at the level of gene transcription. Our findings support the notion that proteostasis decline may have major implications on human aging.

## Introduction

Aging is often characterized by the deterioration in the ability to properly respond to external cues. Various systems that are designed to protect the organism against external assaults lose their ability to efficiently mount a full defense when individuals age (1).

Aging is also accompanied by accumulation of damaged proteins. Damaged misfolded proteins are targets of the Protein Quality Control (PQC system), consisting of the proteasome degradation system, and a network of molecular chaperones, whose role is to identify misfolded proteins, and to either refold them, target them to degradation, or to sequester the damaged proteins (2). The chaperone network is also highly induced in response to a variety of proteotoxic stress conditions that lead to increased load of misfolded proteins, and is often able to efficiently maintain protein homeostasis in the face of fluctuating environments (3). However, damaged proteins which are often efficiently dealt with and cleared in young cells, tend to accumulate with age (4). Accordingly, human diseases of protein misfolding and aggregation, such as Alzheimer’s and Parkinson’s disease, are often considered age-related, and their prevalence increases dramatically with age (1).

Proteostasis collapse has been found and characterized as a hallmark of aging in nematodes (5, 6). Studies have shown that aging in nematodes is accompanied by a decline in the ability of the organism to deal with misfolded proteins, and mount an efficient proteotoxic stress response (5). The proteostasis collapse in nematodes occurs at a specific time frame of adulthood and is linked to the reproductive onset (7–9). Furthermore, a specific chromatin state has been associated with the phenomenon (7).

But while in nematodes, proteostasis collapse in adulthood is well established, whether a similar phenomenon occurs in mammalian species is still unclear. Studies in mammalian organisms over the years have mainly focused on Hsp70, with mixed conclusions; while several studies have found an impaired stress-mediated Hsp70 induction in aging (10–15), as well as impaired DNA binding activity of the HSF1 heat shock transcription factor (15, 16), others have reported that aging has little or no effect on stress-mediated induction of Hsp70 (17, 18).

While the general notion of proteostasis collapse being a part of human aging is plausible, especially given the increased prevalence of proteostasis-related diseases, such as Alzheimer’s, Parkinson’s etc., with age, whether proteostasis decline is in fact a characteristic of human aging still remains unclear. Additionally, there is still debate whether the proteostasis collapse in nematodes is the consequence of damaged proteins that have accumulated throughout the life of the organism, or whether it is a programmed event (2, 6, 9).

Here we decided to directly test whether proteostasis decline is a part of human cellular aging. Cellular senescence is a hallmark of aging; the relationship between cellular senescence and human aging is well established(19), and aged mammalian tissues accumulate senescent cells (20, 21). We therefore chose to use primary human fibroblast cells, which undergo replicative senescence, in order to test the hypothesis of the human proteostasis decline. We exposed young and senescent isogenic cell populations to stress, and further performed an in-depth characterization of the transcriptome and translatome of these cells in response to heat shock using RNA-seq and ribosome footprint profiling. We found that human senescence shows a significant signature of proteostasis decline. In an unbiased analysis of our transcriptome data, a group of more than 160 genes, including many chaperones, showed impaired induction upon heat shock in senescent compared to young cells. Additionally, the nuclear localization of phosphorylated HSF1, the activated form of the heat shock transcription factor, was compromised in senescent cells, as well as its nuclear distribution. Furthermore, alternative splicing regulation, which we found to be prevalent upon heat shock in young cells, is highly diminished in senescent cells, further elaborating the scope of the senescent proteostasis decline. Subsequently, using ribosome footprint profiling, we revealed that the UPR (Unfolded Protein Response) of the ER was activated in young cells, however the coordination between different UPR branches, namely sensing, translation and transcription, was highly impaired in senescent cells; while UPR sensing as well as translational responses were enhanced in senescence, senescent cells were unable to initiate UPR-related transcriptional responses. Finally, we showed that proteasome function is diminished in heat-shocked senescent cells, and could not recover upon return of cells to normal growth temperature. These results further illuminate the broad molecular scope of the proteostasis decline in human cells, pinpointing to specific transcriptional dynamics impairments.

## Results

### Heat shock induction of chaperones is compromised in senescent human cells

In order to directly test the proteostasis collapse hypothesis in human cells, we utilized young primary human WI38 fibroblasts (passage 24), and passaged them until they reached replicative senescence (passage 36 Fig. S1A-C). We synchronized the cells to minimize potential differences due to young cells being in different cell cycle stages (see Methods), and then exposed these young and senescent isogenic cells to 2h of acute heat shock at 44°C.

Following heat shock (HS), young and senescent cells were harvested and RNA-seq was performed to obtain a transcriptome-wide view of gene expression programs (see Methods, and Table S1 for gene expression data). Differential expression analysis (see Methods) identified about 550 genes which were either differentially expressed upon HS, or between young and senescent cells. We further subjected these differentially expressed genes to unsupervised clustering analysis to determine the major behaviors in the data (Fig. 1A, Table S2A). One of the largest clusters resulting from the analysis consisted of 161 genes that were induced in heat shock in both young and senescent cells, however the extent of induction in senescent cells was compromised (Fig. 1A,B). Importantly, the basal mRNA expression levels of the genes in this cluster was similar between young and senescent cells (Fig. 1C). Interestingly, pathway analysis of the genes in this cluster identified stress response genes and chaperones to be highly enriched (Fig. 2A). We further verified this behavior for candidate chaperones using realtime qPCR (Fig. S2A,B). Therefore, this cluster presents the molecular manifestation of a proteostasis decline whereby chaperones are induced during HS in young cells, but their induction is impaired in senescent cells. We thus termed this cluster – the proteostasis decline cluster.

**Figure 1:**
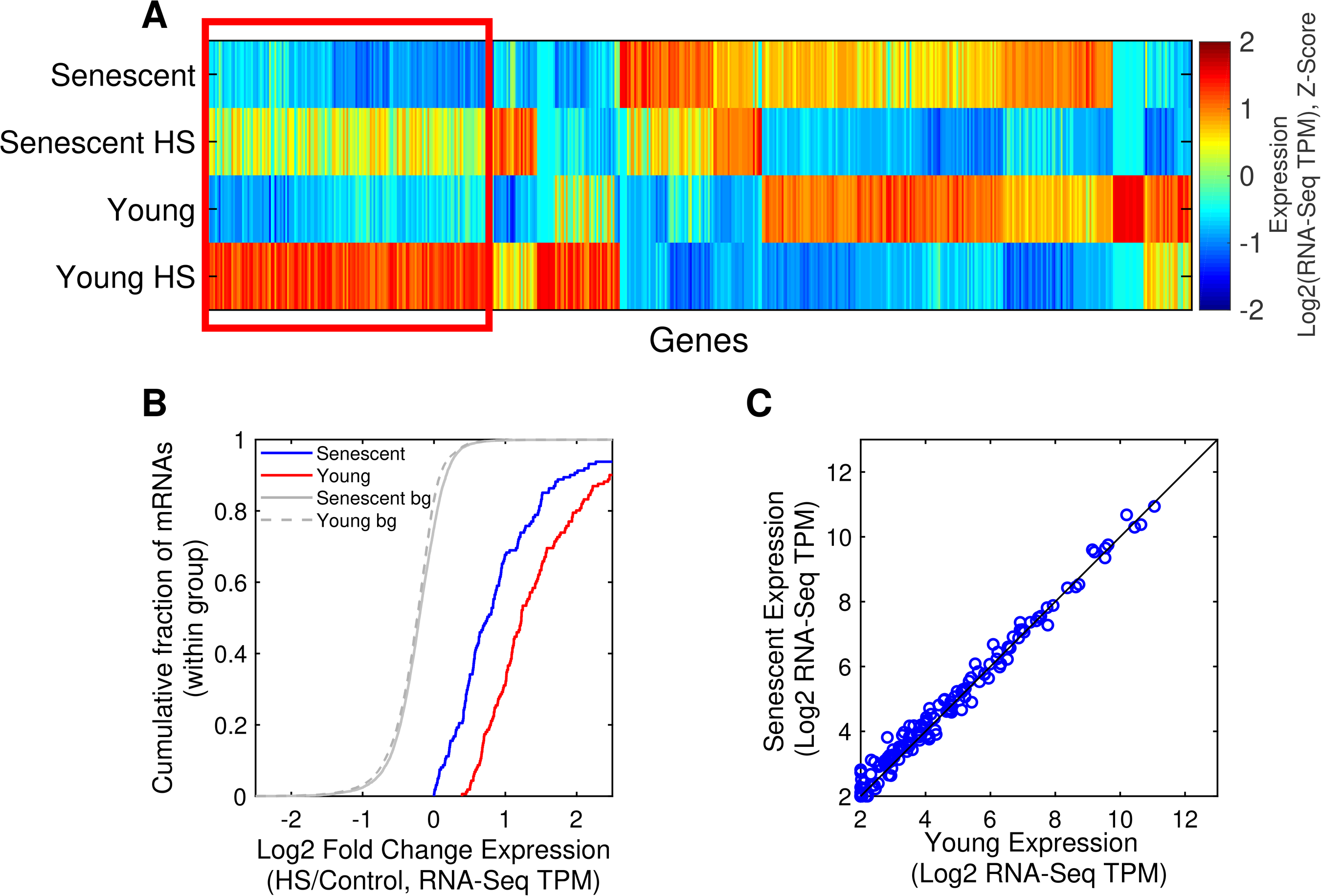
mRNA expression analysis of heat shock treated young and senescent cells. (A) Hierarchical clustering analysis of 548 mRNAs with significant expression difference (using DESeq2 FDR-corrected p-value < 0.05) in at least one comparison between different sample types. The figure shows a gene-wise normalized Z-score heatmap of the log2 expression (RNA-seq TPM) values. The proteostasis decline cluster, a cluster with induced expression levels upon HS, which is attenuated in senescent cells, is marked by a red box. Of the 161 mRNAs in this cluster, there are 27 chaperones, out of 28 differentially expressed chaperones identified. (B) CDF plot of the log2 expression fold change (HS/Control RNA-seq TPM) for mRNAs in the proteostasis decline cluster, shown for senescent (blue) and young (red) cells. Gray lines depict the background distributions (bg), corresponding to all expressed genes in senescent (solid line) and young cells (dashed line). The log2 fold change is greater in young than in senescent (p=4.3^−10^, KS test). (C) No significant difference between basal expression (log2 RNA-Seq TPM) of the mRNAs in the proteostasis decline cluster in young vs. senescent cells (p=0.7, KS test).

**Figure 2:**
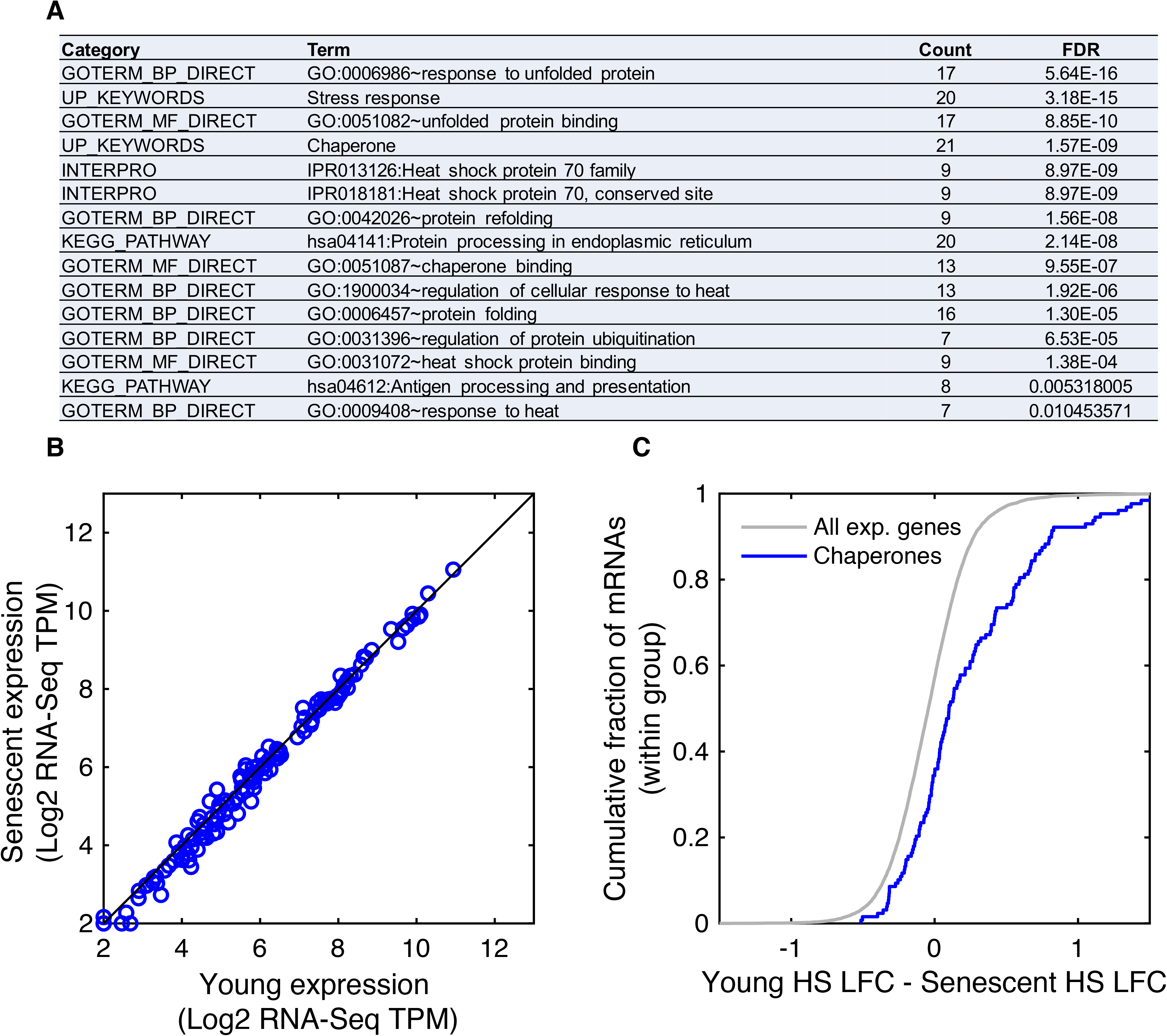
Heat shock-mediated induction of chaperones is impaired in senescent cells. (A) Functional enrichment analysis of the proteostasis decline cluster performed using DAVID, showed that the cluster is characterized by stress response genes and chaperones. (B) No significant difference between mRNA expression levels (log2 RNA-Seq TPM) of all chaperones in young vs. senescent samples (p=1, KS test). Manually curated chaperones list (157 chaperones) is in Table S5, similar results obtained with the chaperone list from Brehme et al. (34), see Fig. S3C-D. (C) CDF plot of the difference between the HS fold changes (log2 TPM HS/Control, denoted as LFC) between young and senescent cells, demonstrating that chaperones are overall more highly induced in young cells (p = 1.4^−9^, KS test).

We note that the opposite behavior, whereby genes were more induced in senescent cells compared to young cells, was not present in the data; a smaller cluster of 24 HS-upregulated genes were similarly induced in young and senescent cells (Fig. S2E,M).

Since chaperones were enriched among the proteostasis decline cluster genes, we turned to look at the expression of all chaperones as a group. Chaperones were not basally differentially expressed between unstressed young and senescent cells (Fig. 2B, S3A). Nevertheless, chaperones overall showed a similar pattern of proteostasis decline, whereby their expression was higher in young cells subjected to HS compared to heat-shocked senescent cells (Fig. S3B-D) and they were more highly induced in young cells in response to HS (Fig. 2C). These trends were evident also when looking at separate chaperone families, with HSP70s, HSP60s and HSP40s all showing a significant proteostasis decline behavior (Fig. S3E-H). These results indicate that the proteostasis decline phenomenon we characterized in human cells is even broader, and spans many of the major chaperone families.

### HSF1 nuclear localization and distribution upon heat shock are impaired in senescent cells

The proteostasis decline observed at the level of transcription, which heavily involved chaperones, suggested that the transcriptional heat shock response (HSR) is compromised in senescent cells. We therefore turned to examine the master regulator of the heat shock response, namely, the HSF1 transcription factor. The overall mRNA levels of HSF1 were not different in young vs. senescent cells (Fig. S4A), and the protein levels were also largely the same (Fig. 3A, S4B,C). HSF1 is known to reside in the cytoplasm under normal conditions, and upon proteotoxic stress it is heavily phosphorylated and translocated to the nucleus, where it induces the transcription of many heat shock genes and chaperones (22). We therefore turned to examine phosphorylated HSF1 localization by immunofluorescence in young and senescent cells. Surprisingly, we observed that while phosphorylated HSF1 gave a strong nuclear staining in young cells treated with HS, in senescent cells there was an additional apparent cytoplasmic staining (Fig. 3B,C, S4E). Non-stressed cells, both young and senescent, showed diffused cytoplasmic staining (Fig. S4D). To further quantify the degree of nuclear localization of phospho-HSF1 upon HS, we performed nuclear-cytoplasmic fractionation in young and senescent HS treated cells, and examined phospho-HSF1 using Western blot (Fig. 3D,E). Non heat-shocked cells showed low cytoplasmic signal for phospho-HSF1, without any nuclear signal. Following heat shock, phospho-HSF1 nuclear levels were high in young cells (Fig. 3D), while in senescent cells the fraction of nuclear phospho-HSF1 was over three-fold lower (Fig. 3D,E). Additionally, we observed that phospho-HSF1 tended to concentrate in a few (1 to 4) distinct nuclear foci in young heat-shocked cells (Fig. 3B,C, S4E), consistent with nuclear stress bodies, as previously characterized in several cell types (23). Senescent cells, however, showed a much more distributed nuclear staining of phospho-HSF1, with significantly higher number of nuclear foci per nucleus, as quantified by image analysis of 3D confocal microscopy images (Fig. S4F,G). Thus, our data suggest that HSF1 nuclear translocation, as well as nuclear distribution, upon heat shock are highly compromised in senescent cells.

**Figure 3:**
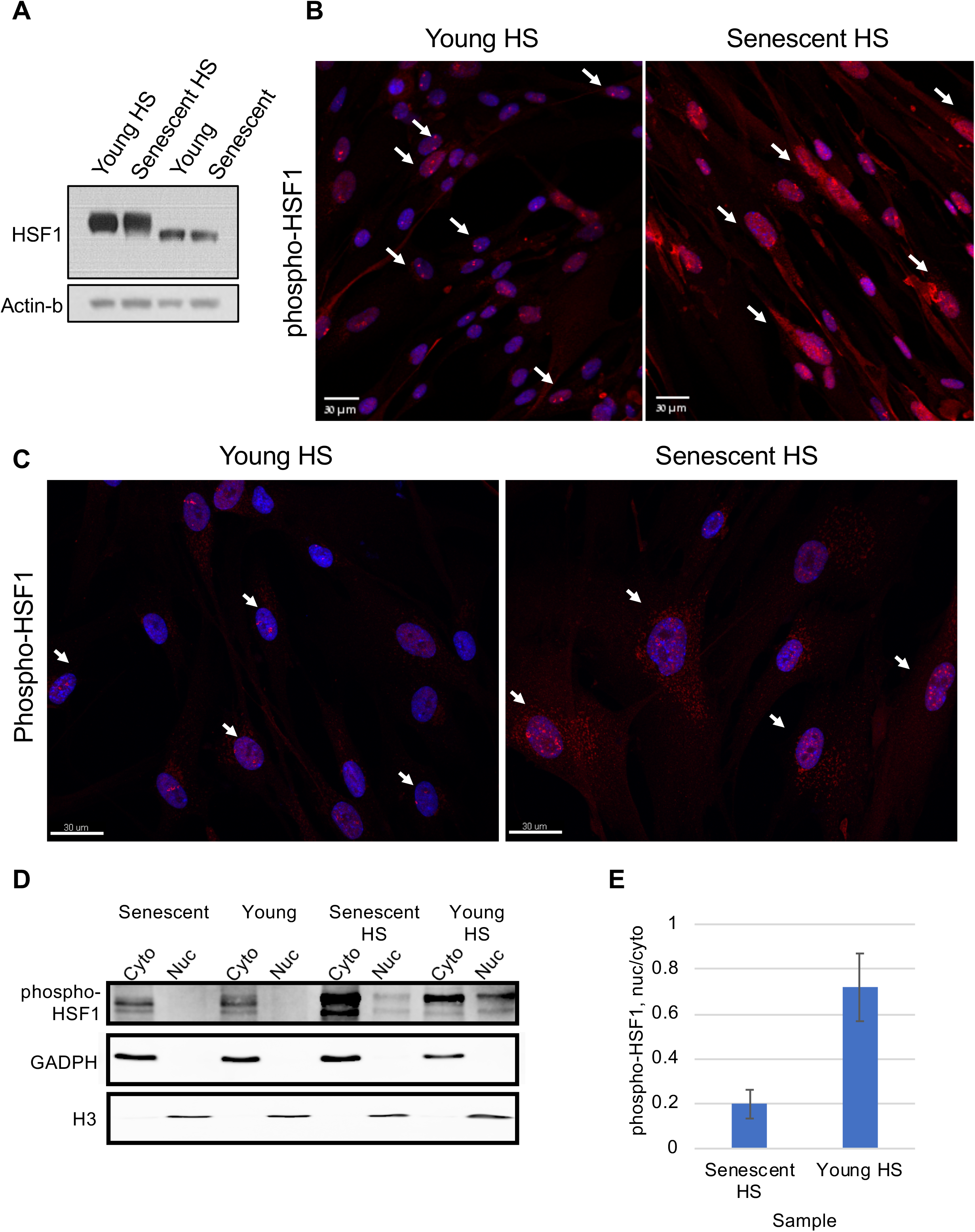
Heat shock-mediated nuclear localization and distribution of HSF1 are hampered in senescent cells. (A) The total levels of HSF1 in young and senescent cells are very similar (see also Fig. S4A-C). (B) Immunofluorescence (IF) staining of phospho-HSF1 in young and senescent heat shocked cells show increased cytoplasmic staining in senescent cells. Additionally, while most young cells show 1-4 single bright nuclear foci of phospho-HSF1 upon HS, most senescent cells show many disorganized foci of phospho-HSF1. White arrows show examples of cells with distinct 1-4 foci in young cells (left) and disorganized HSF1 localization in old cells (right). (C) Confocal 3D imaging of phospho-HSF1 in young and senescent heat-shocked cells revealed a closer look of the nuclear foci distribution in young cells, and its impairment in senescent cells. White arrows show examples of cells with distinct 1-4 foci in young cells (left) and disorganized HSF1 localization in old cells (right). Additional images and quantification of the number of foci is shown in Fig. S4E-G. (D) A representative WB of nucleus-cytoplasm fractionated cells. Phospho-HSF1 is absent from cell nuclei before HS. Following HS in young cells nuclear phospho-HSF1 is increased, whereas in senescent cells the fraction of nuclear phospho-HSF1 is significantly lower. GAPDH and histone H3 present cytoplasmic or nuclear enrichment, respectively. (E) Quantification of WB densitometry for phospho-HSF1, using Fiji. Phospho-HSF1 was quantified in each fraction, and nucleus was divided by the corresponding cytoplasmic fraction. Bars represent mean and std error of three biological replicates.

### Proteostasis decline in senescent human cells is evident at the levels of splicing regulation

We have previously shown that heat shock triggers widespread changes at the level of alternative splicing in mouse cells (24), leading to changes in isoform expression, including extensive yet selective intron retention (24). We therefore decided to explore alternative splicing regulation in the human senescence system. We performed alternative splicing analysis using MISO (25), an algorithm that quantifies the change in the inclusion fraction of a splicing event (Percent Spliced In, PSI), and provides a Bayes Factor (BF) as an estimate for the significance of the change. We analyzed a variety of previously annotated isoform type changes (25), including exon skipping, intron retention, alternative 5’ and 3’ splice site usage, and alternative last exon usage, and quantified the significance of their changes either upon HS in young or senescent cells, or between young and senescent cells. We further defined strict cutoffs in order to enforce replicate reproducibility of the significance of change in alternative splicing (see Methods). Overall, we found 268 significantly changing annotated alternative splicing events in our system (Fig. 4A, S5A-D). There were very few significant changes in alternative isoforms between unstressed young and senescent cells, and the majority of significant alternative splicing changes occurred upon HS (Fig. S5E-J). Surprisingly, we found that, in all alternative splicing categories examined, senescent cells show very few significant splicing changes upon HS, 44-81% less than in young cells (Fig. 4A). We further used clustering analysis to look at the patterns of alternative splicing changes, and observed, again, a clear behavior of proteostasis decline, whereby PSI values show a robust change upon HS in young cells, and changes were much weaker in senescent cells (Fig. S5D). We further examined global intron retention, as in (24), and found that, here too, senescent cells have 70% less retained introns upon HS than young cells (Fig. 4B, S5K). All of these trends were robust to the choice of parameters (see Methods and Fig. S5E-J). Thus, our data suggested that alternative splicing regulation upon HS is largely impaired in senescent cells, further expanding the scope of the molecular proteostasis decline.

**Figure 4:**
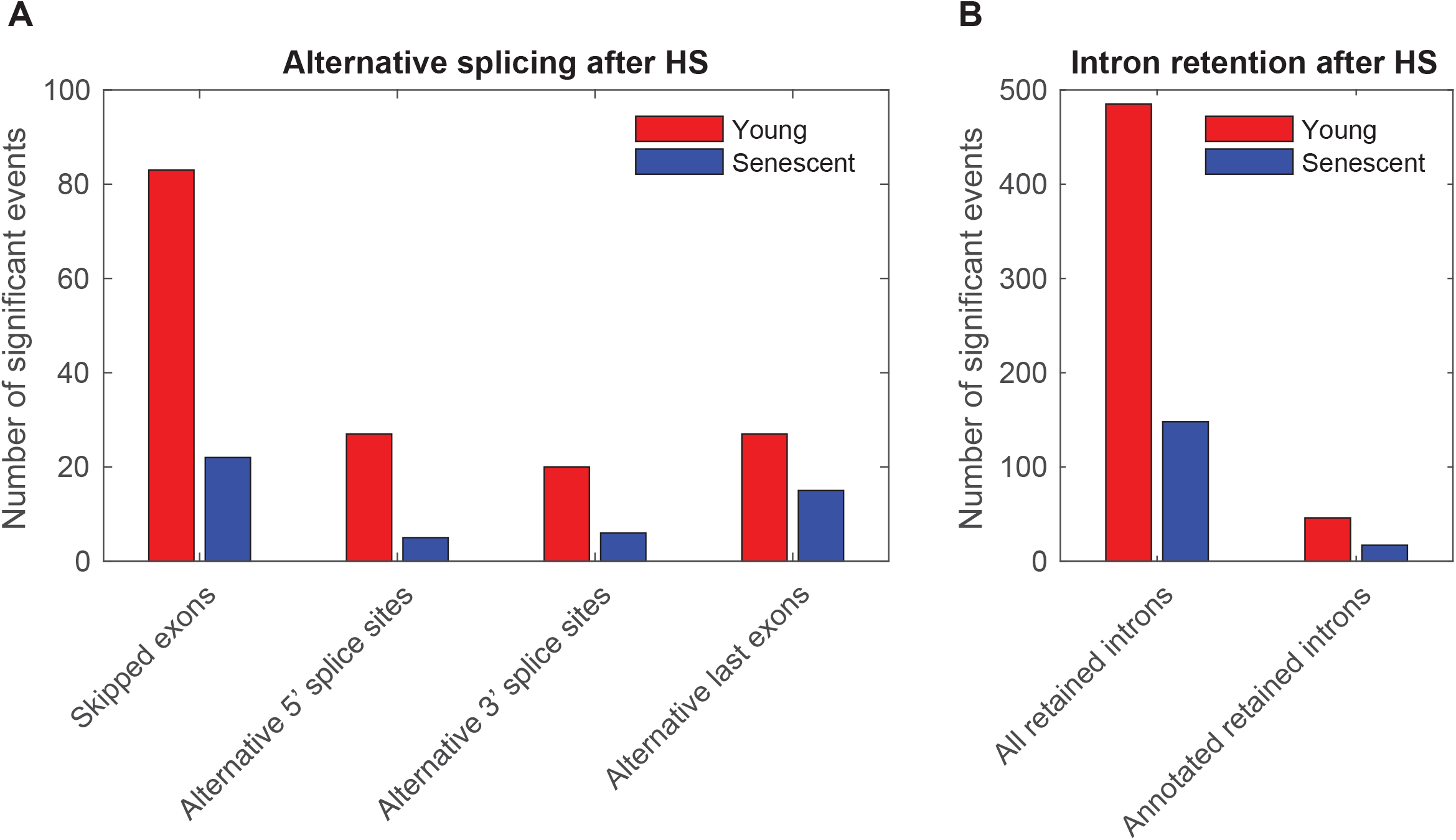
Regulation of alternative splicing following heat shock is highly diminished in senescent cells. Number of differential alternative splicing (A) and intron retention (B) isoforms upon HS in young (red) and senescent (blue) cells. Alternative splicing and annotated retained introns events were initially downloaded from the MISO annotation collection (25). An event was denoted significant if the Bayes Factor (BF) of each HS vs. Control comparisons (in the two replicates) were above eight and the BF of replicate comparisons (Control1 vs. Control2 and HS1 vs. HS2) were below four. The trend is robust to varying BF cutoffs, as shown in Fig. S5E-J.

### Exploring translational control in the young and senescent heat shock response

We next turned to map the cellular translatome in young and senescent cells in response to heat shock, in order to explore potential regulation at the level of translation. To that end, we performed ribosome footprint profiling (or Ribo-seq) on the same young and senescent cells described above, before or after exposing them to 2h of HS (see Methods). Translation level changes largely mirrored mRNA expression changes for the major differentially expressed RNA-seq clusters that we have identified above (Fig. 1A, S2D-K and S6A-H). Differential expression analysis of the translation data resulted in more differentially expressed genes than transcript level data; we found 1222 mRNAs (Fig. 5) with differential translation compared to 548 differentially expressed mRNAs identified by RNA-seq (Fig. 1A). Overall, the two sets of differentially translated or expressed genes partially overlapped (Fig. S6I). Clustering analysis (Table S2) revealed that the proteostasis decline cluster was also evident at the level of translation (Fig. 5, red cluster), and that the translation proteostasis decline cluster highly overlapped with the RNA-seq proteostasis decline cluster (Fig. S6I). Here too, there was no difference in the basal translation levels of the mRNAs in the proteostasis decline cluster between young and senescent cells (Fig. S6N,U). The translation proteostasis decline cluster was enriched with stress response genes and chaperones (Table S3B), similarly to the RNA proteostasis decline cluster.

**Figure 5:**
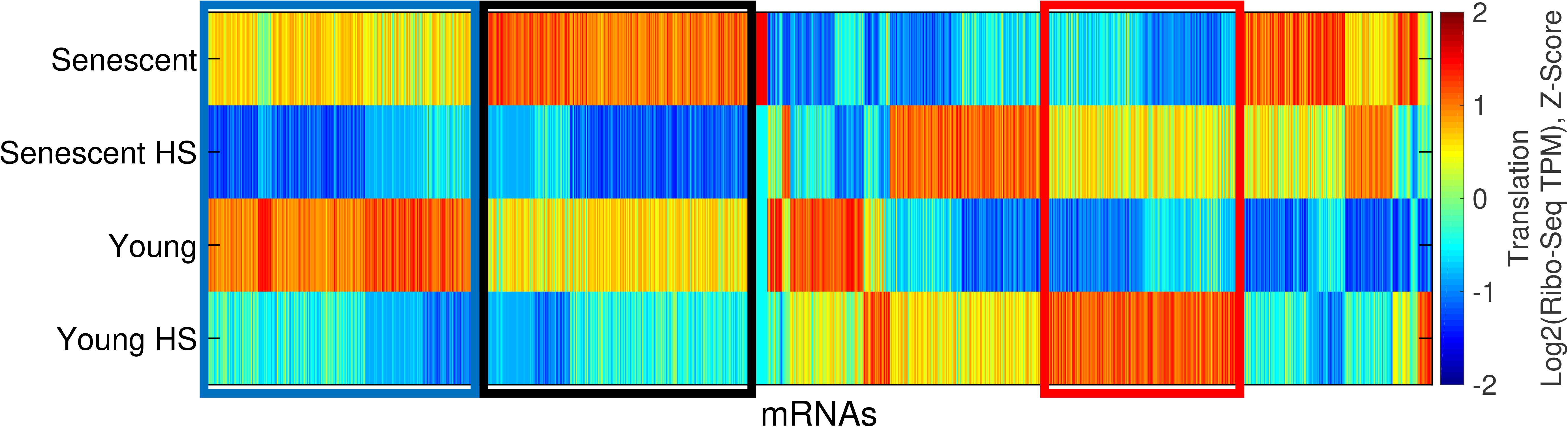
Exploring translational control in the young and senescent heat shock response using ribosome footprint profiling. Hierarchical clustering analysis of 1222 mRNAs with significant expression difference (DESeq2 FDR-corrected p-value < 0.05) in at least one comparison between samples. The figure shows a gene-wise Z-score normalized heatmap of the translation (log2 Ribo-Seq TPM) values. The red box marks a cluster with increased expression level upon HS, which is attenuated in senescent cells, in line with the proteostasis decline mRNA-level cluster above (Fig. 1). Of the 189 genes in this cluster, there are 30 chaperones out of 39 DEG chaperones. The blue and black boxes mark two HS repression clusters, which are both translation specific.

Within the translationally differentially expressed gene group, our clustering analysis has identified a second cluster of HS induced genes (Fig. 5, S6I), with about 150 mRNAs which were HS-induced in senescent cells slightly more than in young cells (Fig. S6AH). These mRNAs were enriched with RNA binding proteins, and with extracellular matrix and extracellular exosome genes (Table S3B). Here too, mRNAs did not show any basal difference in their translation between young and senescent cells (Fig. S6M,T).

### Loss of coordination between UPR regulatory branches in senescent cells

Our clustering analysis of differentially translated mRNAs has identified two large clusters of HS repressed mRNAs (Fig. 5, blue and black clusters). Both of these clusters were largely driven by translational repression (Fig. S6I,X,Y,AE,AF). But while the blue cluster showed similar degrees of repression upon HS in young and senescent cells (Fig. S6AE), the black cluster mRNAs were much more repressed in senescent cells (Fig. 6A). It is therefore denoted as the senescence-enhanced HS repression cluster. Pathway analysis revealed that this cluster is highly enriched with glycoproteins, disulfide bond containing proteins, and membrane proteins (Fig. 6B, Table S4), which represent the major classes of ER targets. Widespread repression of ER targets has been previously reported by us and others to be a hallmark of the UPR (26–28), representing a significant translational regulatory response mediated by PERK (26). We therefore reasoned that the HS treatment in our experiment has induced the UPR. Indeed, examination of the set of mRNAs that were previously identified as a PERK-dependent translational repression UPR signature (26), which was found to be highly enriched for ER targets, confirmed that this set was significantly repressed in both young and senescent cells in response to HS, however repression was substantially greater in senescent cells (Fig. S7A,B). Importantly, here too, repression occurred at the level of translation (Fig. S7A,B). Western blot analysis for PERK showed a phosphorylation-related upshift in both young and senescent heat-shocked cells (Fig. S7C). Previously, we have shown that the PERK-mediated repression of ER targets as part of the UPR involved eIF2α phosphorylation (26). We confirmed that eIF2α phosphorylation occurred in both young and senescent cells upon HS, to a similar extent (Fig. S7D,E). ATF4 ORF translation, another UPR hallmark event downstream of PERK and phospho-eIF2α (29), was induced to a similar extent in young and senescent heat-shocked cells (Fig. S7F-G). Therefore, it seems that the PERK branch of the UPR is activated in both young and senescent stressed cells. We thus sought to examine other UPR branches in this system.

**Figure 6:**
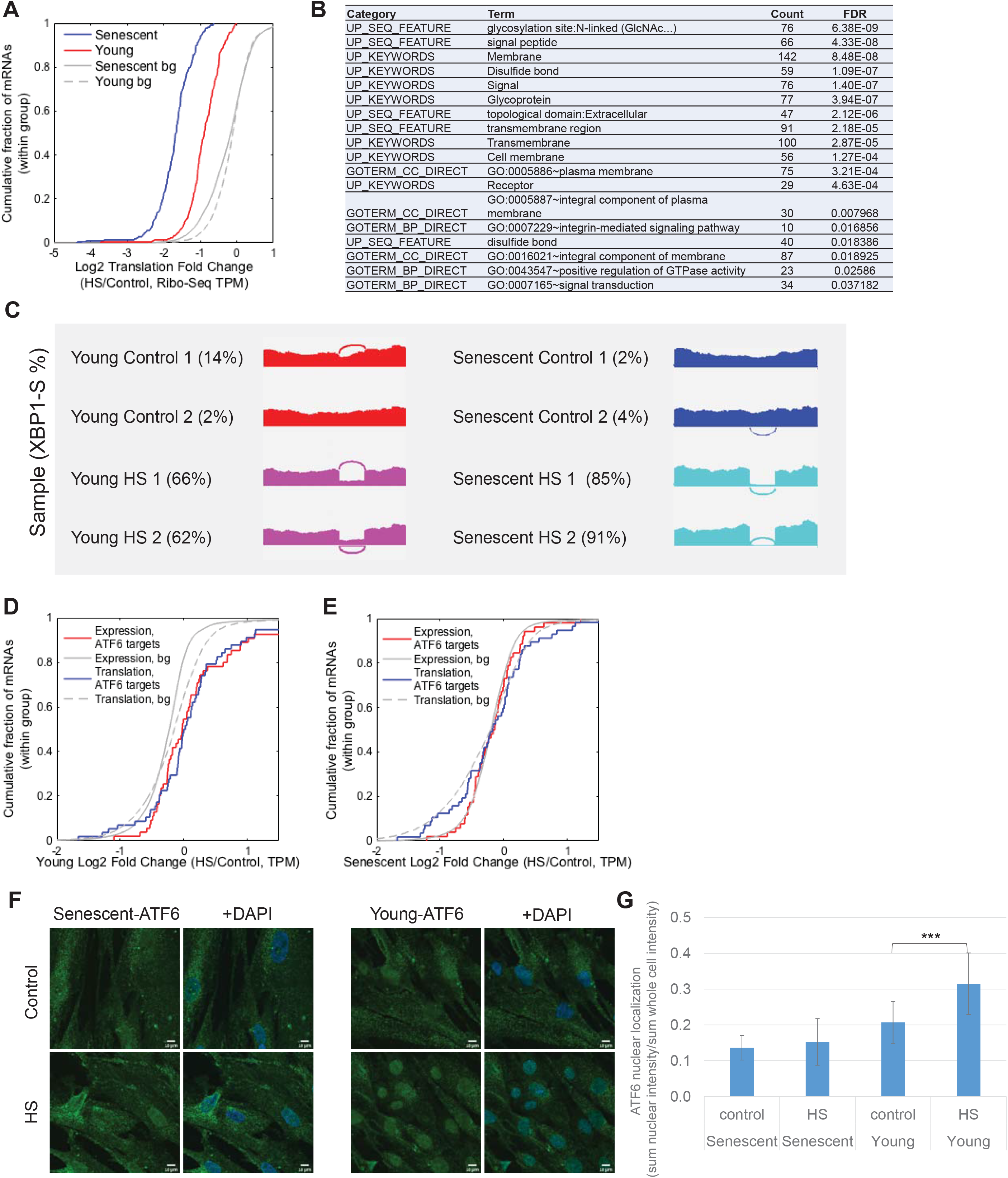
Loss of coordination between UPR branches in senescent cells upon heat shock. (A) CDF plot of the log2 translation fold-changes (HS/Control, Ribo-seq TPM) for mRNAs from the senescence enhanced HS repression cluster (Fig. 5, black cluster), shown for senescent (blue) and young (red) cells. Gray lines depict the corresponding CDF plots for the background distribution, i.e. all translated mRNAs in senescent (solid grey line), or young (grey dashed line) cells. The repression is significantly stronger in senescent than young cells (p=5.9^−44^, KS test). (B)Functional enrichment analysis of the senescence enhanced HS repression cluster was performed using DAVID, and showed that the cluster is highly enriched with ER targets. Full list of annotations is available in Table S4. (C) Splicing plot (Sashimi plot (25)) of XBP1 exon four (of the unspliced isoform). The percent spliced isoform, PSI values, of the spliced XBP1 isoform for all samples was quantified using MISO (25) are indicated: 2%, 4% (senescent) 85%, 91% (senescent HS) 2%, 14% (young) 66%, 62% (young HS). Significance of change was quantified by MISO and resulted in the following Bayes factors (BFs): BFs for HS vs. Control samples: 1012, for all comparisons. BFs for senescent vs. young: 1.9,0.1, indicating no basal difference in the amount of XBP1-spliced. BFs for senescent-HS vs. young-HS: 14717,3.810. (D, E) CDF plots of the log2 fold change (HS/Control) of the set of bona-fide ATF6 target genes (taken from (31)) demonstrate their significant induction upon HS in young cells (D) both at the mRNA (red) and the translation (blue) levels (p=1.7^−5^, and p=1.1^−3^ for mRNA expression and translation respectively, using KS test). No induction was observed in senescent cells (E, p=0.87 and p=0.77 for mRNA expression and translation respectively, using KS test). (F) ATF6 immunofluorescence (IF) confocal microscopy imaging showed increased nuclear localization in young heat shocked cells, with a diminished trend in senescence. DAPI marks nuclei. Additional fields are shown in Fig. S7K-AH. (G) Image analysis was performed on IF images. Mean and std of ATF6 nuclear localization are presented, calculated as the sum of fluorescence intensity in the nuclei divided by the sum of fluorescence intensity in whole cells (see Methods), with a total of 16,17,19 and 20 images taken for Young, Young HS, Senescent and Senescent HS samples respectively, from three biological replicate experiments. Significance was calculated using t-test, ***=2*10^−4^. Data for the three separate replicates is shown in Fig. S7AI-AJ.

To further explore the induction of the IRE1 branch of the UPR, we examined the splicing profile of XBP1, whose cytoplasmic splicing by IRE1 is considered to be one of the primary events in ER stress sensing (29). To that end, we quantified the degree of splicing in each sample using the MISO algorithm (25). First, there was very little XBP1 splicing in both young and senescent unstressed cells, verifying that senescent cells are not under basal ER stress. Upon heat stress in young cells, XBP1 is partially spliced, to a level of about 64% (Fig. 6C, S7H). In senescent cells, however, XBP1 splicing was enhanced, to a level of about 88% (Fig. 6C).

Therefore, it seems that senescent cells sense protein misfolding in the ER that occurs in heat shock to an even higher degree than young cells. They also activate the PERK branch and its translational outputs, both translational enhancement (ATF4, Fig. S7F,G) and repression (ER targets, Fig. 6A,B, Fig. S7A,B), to a similar, or even higher extent than in young cells. Nevertheless, in order to transition into the adaptive phase of the UPR, transcriptional activation of the target genes of the two transcriptional UPR branches, namely the ATF6 and XBP1-s (XBP1-spliced) transcription factors (30), should take place. We therefore examined the transcriptional outputs of these two branches, by looking at a set of bona-fide ATF6 and XBP1-s target genes previously defined by Shoulders et al. (31). Strikingly, we found that while ATF6 target genes were significantly induced in young cells both transcriptionally and translationally (Fig. 6D, p=1.7^−5^ and p=1.1^−3^ at the mRNA expression and translation levels respectively), they were not induced at all in senescent cells (Fig. 6E, p=0.87 and p=0.77 at the expression and translation levels respectively). A similar trend emerged when we examined the set of bona-fide XBP1-s targets (31) (Fig. S7I,J).

Interestingly, further examination of ATF6 localization using immunofluorescence imaging and image analysis revealed that, in young cells, ATF6 nuclear localization at 2h of heat shock was significantly increased (Fig. 6F,G, S7K-AJ, see Methods). Senescent cells, however, did not show a consistent increase in ATF6 nuclear localization following heat shock (Fig. 6F,G, S7K-AJ).

Hence, our data suggest that while young cells are able to sense ER protein misfolding upon HS, and initiate both transcriptional activation as well as translational activation and repression programs as part of the UPR, senescent cells show decoupling of the different UPR branches; Their IRE1-mediated stress sensing (as quantified by XBP1 splicing), and PERK-mediated translational regulation are both intact, or even enhanced, however they fail to activate the UPR transcriptional branches of ATF6 and XBP1-s. Thus, our combined analysis of transcription and translation data unraveled yet another level of proteostasis decline in senescent cells. Furthermore, our data revealed what seems to be a more general deterioration in the ability of senescent cells to initiate the required transcriptional reprogramming upon protein misfolding stress, by HSF1, ATF6 and XBP1-s, an essential step in the cellular ability to cope with and adapt to stress.

### Deterioration of proteasome function in stressed senescent cells

Finally, we asked how other aspects of protein homeostasis regulation might be affected by heat stress in senescent cells. Specifically, we examined the proteasome, as an essential part of the PQC system responsible for degradation and clearance of misfolded proteins (32). Previous studies reported that late passage WI-38 cells showed lower levels of several proteasome beta subunits, and this was accompanied by a reduction in basal proteasome activity (33). In our system, however, where cells have just fully entered senescence, the levels of proteasome components were unchanged relative to young cells (Fig. 7A,B, S8A-D); mRNA levels and translation levels of all proteasome subunits was unchanged between young and senescent cells (Fig. 7A,B, S8A,B), and protein levels of the beta2 subunit by WB were similar as well (Fig. S8C,D). We next measured proteasome activity (as in (33), see Methods), and found that basal proteasome activity was similar between young and senescent cells (Fig. S8E). Strikingly, however, while at 2 hours of heat shock proteasome activity in young cells remained similar to that of unstressed cells, senescent cells showed a significant reduction of over 30% in proteasome activity following heat shock (Fig. 7C). Moreover, senescent cells were unable to restore their proteasome activity after 4h of recovery post heat shock (Fig. 7C), with a 45% reduction compared to unstressed cells. This is in sharp contrast to young cells, that had intact proteasome function also after recovery (Fig. 7C).

**Figure 7:**
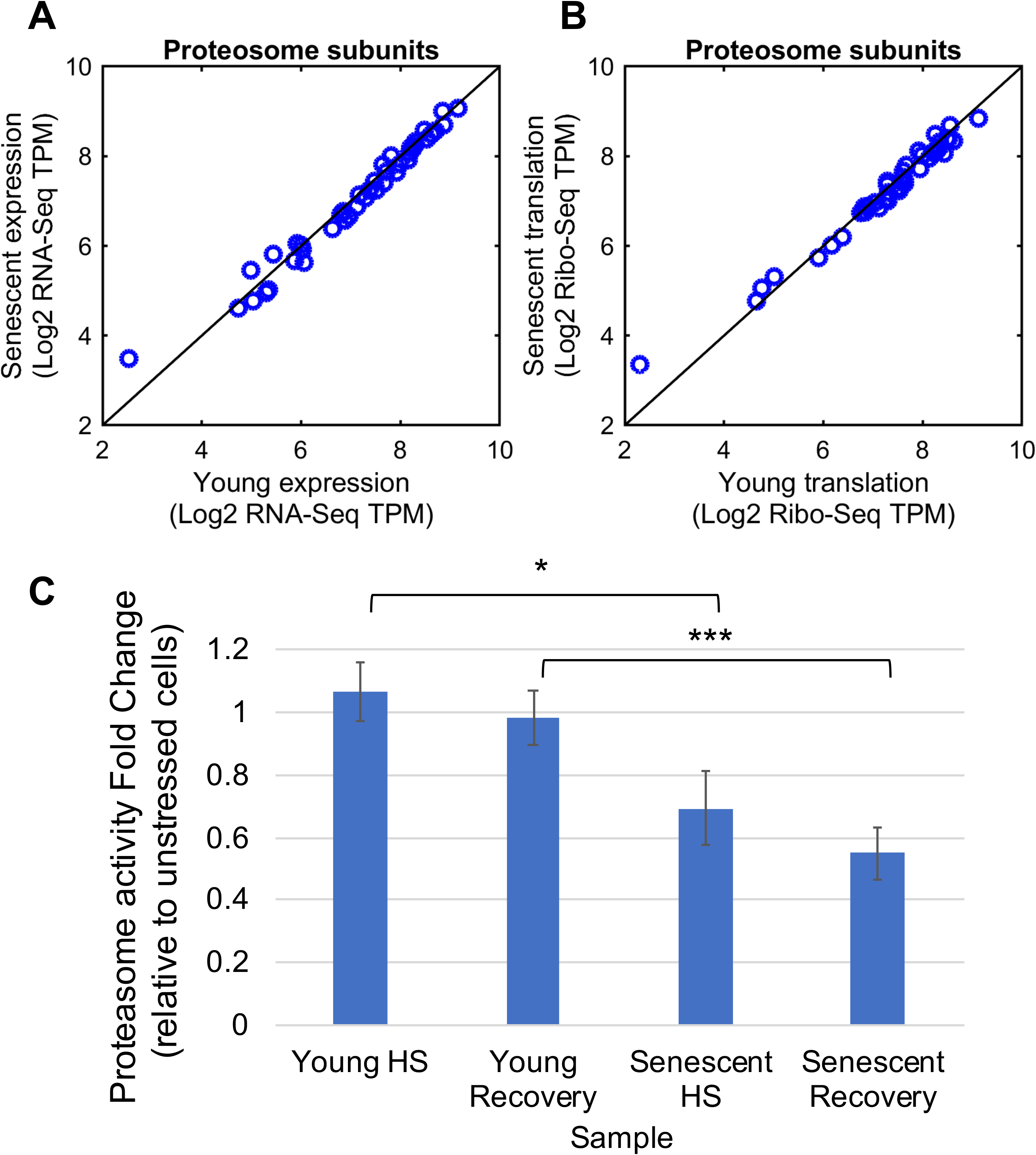
Deterioration of proteasome function in stressed senescent cells. (A-B) Proteasome subunits show no difference in mRNA levels (A) or translation levels (B) between young and senescent cells, as evident from RNA-seq (A) and Ribo-seq data (B). This holds true also when comparing young HS vs. senescent HS cells (Fig. S8A-B). (C) Proteasome activity assay was performed as in (33) using the Suc-LLVY-AMC fluorogenic proteasome substrate (see Methods). The assay was performed on lysates of young and senescent cells, either untreated, after 2h of HS, or following 2h of HS and a subsequent recovery of 4h at 37°C. Activity of all samples was subtracted with MG132 treated lysates for background, and then HS and recovery samples were divided by untreated cell values to assess the change in proteasome activity. While in young cells proteasome activity is unchanged during stress and recovery, senescent cells show significantly diminished activity of 31% and 45% in HS and recovery respectively (t-test p-values are shown, *=0.03, ***=0.005). Mean and std error are presented of 5-7 biological replicates.

Thus, the human senescence proteostasis decline is not only manifested in deteriorated transcriptional responses, but also has implications on the cellular proteasome function in both stress and recovery.

## Discussion

An impaired cellular stress response during aging has long been thought to contribute to age-related diseases (2, 20). In nematodes, the phenomenon of proteostasis collapse upon aging is well established (5), and has been shown to initiate at the onset of reproduction (7, 8). Furthermore, multiple lines of evidence suggest that it is a programmed event, related to overall organismal reproductive capacity and fitness (2, 8). Here, we report the first genome-wide quantification of transcriptional and translational responses to heat shock in senescent human cells. Our results show that proteostasis decline occurs in human primary fibroblasts upon replicative senescence. It is a widespread phenomenon including a reduced transcriptional induction of many chaperones and impairment of splicing regulation, as well as a diminished proteasome function following stress.

Proteostasis collapse in nematodes has been shown to represent an organismal level phenomenon. Furthermore, the nature, extent, and onset of the nematode proteostasis collapse can be modulated by external signals in a non-autonomous fashion (2, 6), however whether or not it naturally occurs in a cell-autonomous fashion has remained unknown. Our data here showed that the proteostasis decline in human senescence occurs in a cell-autonomous manner, namely, it is an intrinsic characteristic of cells upon entry to a senescent state. These findings further support the notion that this is a programmed phenomenon, rather than a mere consequence of accumulation of damaged proteins throughout organismal life. The question of the consequences of the human cellular senescence proteostasis decline at the organismal level is very intriguing, although highly complex. The connection between human aging and senescence has been well established for many years (19). Our data therefore imply that indeed, accumulating population of senescent cells in the aged human organism experience proteostasis decline. To examine the potential link between our findings and human aging, we analyzed a dataset of chaperone signatures previously identified as consistently altered in aged human brains (34). Interestingly, we found that while the levels of both aging brain-induced and -repressed chaperones were similar between young and senescent cells (Fig. 8, S9), chaperones identified as repressed in aging brains, but not those that were induced, showed the proteostasis decline behavior in senescent human cells; namely, they were significantly more induced upon heat shock in young cells than in senescent cells (p=8.5^−7^ – 7.3^−4^, Fig. 8A-C, S9A-C). Potentially, diminished steady state expression of a subset of chaperones in aged brains is further accompanied by the inability to induce these chaperones upon proteostasis insults. This could be the manifestation of a combined overall effect of programmed proteostasis decline, as well as proteostasis deterioration due to gradual accumulation of damaged proteins that burden the chaperone network upon human aging. Therefore, our study represents a first step towards generalizing the understanding of proteostasis decline in human aging, and future work will further delineate the cell-autonomous vs. non-autonomous nature of the phenomenon in aged mammalian organisms.

**Figure 8:**
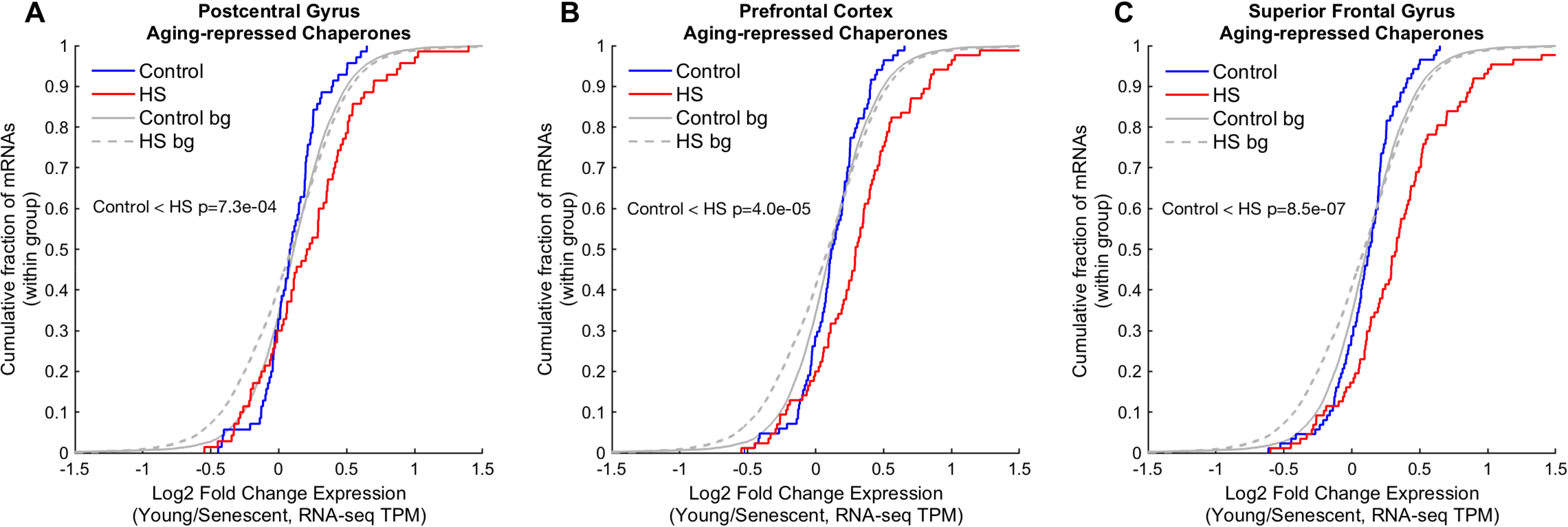
Repressed chaperone signature from aged human brains show proteostasis decline behavior in human senescence. (A-C) Age-repressed chaperones were taken from Brehme et al. (34), as defined for three different brain regions. CDF plots depict the log2 expression fold changes of this signature in Young/Senescent, either in untreated (blue) or heat-shocked (red) cells. These chaperones show a proteostasis decline behavior in human senescent cells: they were significantly more induced in heat shock in young vs. senescent cells, and therefore the HS curve is significantly shifted. On the other hand, aged-induced chaperones from the same tissues show no significant difference (see Fig. S9).

Our evidence point to stress sensing being intact in senescent cells, as shown by greater activation of the IRE1-mediated XBP1-splicing branch of the UPR (Fig. 6C,S7H). Additionally, we observed an intact translational regulatory response, evident by enhanced activation of the PERK-mediated UPR translation signature (Fig. 6A,B,S7A,B), as well as PERK phosphorylation (Fig. S7C), eIF2α phosphorylation (Fig. S7D,E), and ATF4 ORF translation (Fig. S7F,G). As primary fibroblasts secrete many extracellular matrix proteins, they are apparently more sensitive to various forms of proteotoxicity, including HS, and therefore activated the UPR. Importantly, unstressed senescent cells did not show basal UPR activation, as evident by similar expression levels of BiP (Hspa5, Table S1), and low levels of XBP1 splicing (Fig. 6C). Furthermore, senescent cells in general, and senescent fibroblasts in particular, are highly secretory (35), which could explain why their UPR sensing and translational responses are both aggravated in response to HS. However, eliciting transcriptional responses was compromised in senescent cells not only for HSF1, but also for the UPR transcriptional branches mediated by ATF6 and XBP1-s. Furthermore, our data showed that nuclear localization upon heat stress was impaired for both HSF1 and ATF6. Interestingly, nematode UPR activation is also deteriorated during aging (5, 36), however in contrast to human senescence, nematodes fail to activate the IRE-1 branch altogether (36). In the case of HSF1, nuclear localization in nematodes was actually intact (7). Our evidence suggest that senescent human cells can initiate the UPR, however their ability to transition to the adaptive stage of the UPR, as well as the HSR, where transcriptional programs need to take place, is diminished. Taken together, these evidences suggest that perhaps certain properties required for dynamic stress transcriptional responses are overall declined in senescence. What could be the underlying mechanism for a general deterioration in the ability to elicit stress transcriptional responses in senescent cells? Our data point to two potential, not mutually exclusive, possibilities discussed below.

Previous studies have shown diminished DNA binding activity of HSF1 in human senescence (15). Additionally, HDAC1 has been shown to attenuate HSF1 DNA binding activity upon heat shock in senescent MEFs (16). Our data adds two additional dimensions to explain HSF1 deteriorated transcriptional activity: reduced nuclear localization, and disordered nuclear distribution.

An interesting possibility is that chromatin structure in human senescence in particular, and during aging in general, is altered in a manner that disables efficient dynamic changes in opening of chromatin regions, either globally, or specifically at regions essential for stress responses, thereby affecting access of HSF1, ATF6, and other stress transcription factors to their DNA targets upon stress. In nematodes, proteostasis collapse was accompanied by increased H3K27me3 chromatin marks at several chaperone gene loci (7). Additionally, the expression of the histone demethylase *jmjd-3.1* is decreased in aged nematodes, while its overexpression maintained proper induction of the HSR upon aging (7), and led to increased lifespan (37). Similar trends of reduction in activating histone marks and induction of repressive marks have been shown in flies (38) and fish (39), supporting the notion that a closed chromatin state at chaperone gene loci, or perhaps more globally, may confound the deteriorated induction of the HSR upon aging.

In mammals, however, characterization of senescence and aged chromatin states has yielded a highly complex picture. The chromatin landscape of senescent cells seems to be very different than the pre-senescent state (40). While Senescence-Associated Heterochromatin Foci (SAHF), regions of highly dense heterochromatin, were shown to be a hallmark of human senescence (40, 41), other studies reported a general decrease in histone content in senescent and aged cells (42, 43), and some histone variants were shown to be upregulated (44). Reorganization in the landscape of several activating and repressing histone marks has also been observed (40). Finally, transcription of satellite repeats has been shown to occur after cells enter senescence (45). Interestingly, HSF1 localization to nuclear foci upon heat shock, observed in young cells (Fig. 3B,C,S4E), has been previously demonstrated to occur at satellite III regions (46). Our observation of disorganized HSF1 nuclear foci might be explained by an altered chromatin state which allows more pervasive satellite repeat expression prior to stress, thus leading to redistribution of HSF1 into numerous foci in senescent cell.

Nuclear Lamin disfunction has also been linked to aging and senescence; Lamin B shows decreased expression upon senescence (47, 48), with major consequences on nuclear and genome organization (49), and Lamin A/C mutations causing premature aging in HGPS alter nuclear structure, transcriptional deregulation, and chromatin organization (50, 51). More recently, senescent cells were shown to contain cytoplasmic blebs of chromatin (42, 52). Our results on HSF1 and ATF6 diminished nuclear localization fit well with the notion of compromised nuclear integrity in senescence. If nuclear integrity is compromised, together with reorganized chromatin that disfavors optimal stress transcription factor binding, HSF1, ATF6, and potentially other transcription factors, will remain unbound to chromatin and therefore might more easily exit back into the cytoplasm.

Future studies will unravel the full extent of nuclear integrity deterioration upon senescence, and the link between the altered chromatin landscape to the deterioration in dynamic stress response activation in human senescence and aging.

## Materials and Methods

### Cell culture and stress

WI38 human lung fibroblasts were grown in MEM Earl’s medium, 10% serum, L-glutamine, 1% Na-pyruvate, 1% nonessential amino acids and 1% penicillin/streptomycin solution. For the synchronization of the cell cycle, cells were plated on 15 cm plates at a confluency of 1.5×10^6^ for senescent or 1.3×10^6^ for young cells. The next day, the cells underwent serum starvation (starvation medium was MEM Earl’s medium +1% penicillin/streptomycin solution) for 24 h followed by recovery in regular growth media for another 24 h. After recovery cells were subjected to stress, (heat shock at 44^0^C, for 2 h) and then harvested in Lysis Buffer (20mM Hepes pH 7, 100mM KCl, 5mM MgCl2, 0.5% Na DOC, 0.5% NP-40, 1mM DTT, Protease inhibitors (Roche)). Samples were used for RNA-Seq or Ribo-Seq.

### RNA-Seq library preparation

RNA was isolated from collected samples using the TRIZOL reagent (Thermo) following treatment with Turbo DNase and phenol chloroform precipitation. After quality control check, the samples were subjected to Illumina kit (TruSeq RNA) for library preparation.

### Ribosome footprint profiling library preparation

Ribosome footprint profiling was performed as described in (53) with the following modifications: Following heat shock, cells were lysed directly on the plate on ice for 10 minutes using the Lysis Buffer (see above) without Cycloheximide. Then the samples were treated with 20μl Turbo DNase for 5 minutes at 25^0^C while rotating. At this point 10% of the lysate was taken for RNA-seq (see above). Nuclei were removed with 10min centrifugation at maximum speed at 4^0^C and supernatant was retained. Then, 8 μl NEB RNase If (NEB M0243S) per OD260 of ~10 was added in 2ml of lysis buffer, and was rotated for 55min at room temperature. Ribosomes were pelleted using a sucrose cushion (10ml lysis buffer layered on top of 12.5 ml cushion: 20 mM HEPES pH 7, 100 mM KCl, 5 mM MgCl2, 0.5 M Sucrose), by ultracentrifugation at 60K RPM for 1 hour and 50 minutes, at 4^0^C with a Ti70 rotor.

### RNA-Seq analysis

RNA-Seq reads were filtered for rRNAs, tRNAs, microRNAs, and snoRNAs using STAR (54). The remaining reads were mapped to hg19 version of the human genome using STAR (54). Expression levels were quantified using RSEM (55) to produce gene level quantification TPM (Transcript Per Million) values. TPM values were then averaged between sample replicates.

### Ribosome profiling analysis

Ribosome footprint reads were trimmed to a maximal length of 34 and polyA sequences which were generated in the polyA tailing step of the ribosome footprint profiling protocol were removed, such that footprints of lengths 22-34 were considered. Reads were filtered for rRNAs, tRNAs, microRNAs, and snoRNAs using STAR (54). The remaining reads were mapped to hg19 version of the genome using RefSeq modified CDSs; Transcripts shorter than 100 nucleotides were filtered out and 30 nucleotides were clipped from the start and end of each CDS before mapping, similarly to Ingolia (56). Expression levels (mRNA TPM) were quantified using RSEM (55) after mapping to clipped CDSs using Bowtie2 (57). TPM values were averaged between sample replicates.

### Differential expression analysis

We applied the R package DESeq2 (58) on the read-count tables (rsem.genes.expected_count.txt files) resulting from RSEM to identify mRNAs that were differentially expressed (DEGs) between pairs of samples (we used FDR-corrected p-value < 0.05). We compared either between young and senescent, or between either young or senescent cells before and after heat shock treatments, and all DEGs found in at least one comparison were considered for further analysis.

### Hierarchical clustering

Lowly expressed mRNAs with TPM value below four across all samples were filtered out. TPM values of all remaining genes were thresholded to four. Hierarchical clustering (using MATLAB) of gene expression levels was done based on spearman correlation between log2 TPM values of mRNAs across samples. Clustergrams were displayed after gene-wise Z-score normalization for visualization purposes.

### Functional Enrichment analysis and gene groups

Pathway enrichment analyses were conducted using RDAVIDWebService R package (59), using all expressed genes as background. Pathways with Benjamini corrected p-value <0.05 were designated as significantly enriched.

Chaperone list (Fig. 2B,C, S3A,B) was manually curated (Table S5), or taken from Brehme et al. supplementary information (34) (Fig. S3C-D), for all cytosolic chaperones (excluding the ER and MITO families), and excluding the TPR, PFD families. Additionally, chaperone families HSP70, HSP40 and HSP60 (Fig. S3E-H) were taken from Brehme et al. (34) supplementary information. XBP1 and ATF6 targets were defined by Shoulders et al. (31) following specific direct activation of XBP1 or ATF6 (termed XBP1 and ATF6 targets). PERK-dependent UPR repression signature gene set was defined according to Gonen et al. (26). Groups of chaperones significantly altered (induced or repressed) in aged brains (in Postcentral Gyrus, Prefrontal Cortex and Superior Frontal Gyrus, Fig. 8, S9) were taken from Brehme et al. (34) supplementary information.

For all gene group comparisons using CDF plots, two-sample Kolmogorov-Smirnov goodness-of-fit test (KS test) was performed for every group compared to the corresponding background distribution, using all expressed genes as background, and p-values <0.01 were designated as significant. In some cases, the same test was applied between two groups of interest (indicated). Significant p-values are indicated on the top of the CDF plots, or in the figure legends.

### Nuclear-cytoplasmic fractionation and WB analysis

In order to separate between nuclear and cytoplasmic fractions we used NE-PER Nuclear and Cytoplasmic Extraction Reagents (Thermo Scientific) or hypotonic fractionation protocol (see below). Cells were plated on 6-well plates and after 2h HS were collected in PBS, centrifuged and cell pellets were frozen. The fractionation procedure was performed following manufacturer’s instructions for the NE-PER kit. For hypotonic fractionation, cell were lysed in hypotonic lysis buffer (20mM HEPES, 10mM KCl, 2mM MgCl2, 1mM EDTA, 1mM EGTA, 1mM DTT with protease inhibitors (Roche, 11836170001) and phosphatase inhibitors (Sigma, P5726)), incubated on ice for 15min, then passed 10 times through a 25G syringe, and left on ice for 20min. Cytoplasm was separated from nuclei by centrifugation at 15000g for 5min at 4^0^C. Pellets (nuclei) were washed once with cold PBSx1, then centrifuged again and resuspended in TBS with 0.1%SDS, and sonicated in an Ultrasonic Cell Disruptor (cap tips) at 100% amplitude for 5sec 3 times. For both protocols, protein concentration of cytoplasmic fraction was measured using the BCA kit (Pierce, catalog # 23227) and equal cytoplasmic amounts were loaded. Nuclei amounts were scaled to their respective cytoplasms. At the end, samples were subjected to WB with anti phospho-HSF1 (S326) antibody (Abcam, AB-ab76076), anti H3 (Abcam ab1791) and anti GAPDH (Abcam ab8245).

Additional antibodies used for WB: anti phospho-eIF2α (Cell Signaling, CST-9721S) and anti eIF2α (Cell Signaling, CST-5324S) in Fig. S7D,E, anti HSF1 (10H8 clone, Stressmarq, SMC-118D or Cell Signaling CST-4356s) in Fig. 3A and S4B,C, anti PERK (Santa Cruz, sc-377400) in Fig. S7C, and anti beta2 (Santa Cruz, sc-58410) in Fig. S8C,D. WB bands densitometry were quantified using Fiji.

### HSF1 immunofluorescence

For Immunofluorescence (IF), young or senescent cells were plated on coverslips in 12-well plates. On the day of the experiment, the cells were subjected to a 2h HS and then were fixed in 4% PFA. The antibodies used were: anti phospho-HSF1 (S326) antibody (Abcam, AB-ab76076) at 1:200 dilution, secondary anti-rabbit antibody (Jackson Laboratories) at 1:400 dilution. For the detection of nuclei the coverslips were incubated with 1 ug/ml DAPI in PBS for 5 min before mounting. Images were taken with Zeiss LSM700 laser scanning confocal microscope. Image analysis for quantification of the number of phospho-HSF1 nuclear foci was performed using Imaris software scripts on 3D confocal microscopy images (x40 magnification), with 6-8 fields for each of either young HS or senescent HS cells. Two independent replicate experiments gave similar results (Fig. S4F,G).

### ATF6 immunofluorescence and image analysis

Young or senescent cells were seeded at 12k and 15k cells/well respectively in 8-chambered μ-Slide (ibidi, 80826) or 25k and 32k cells/well in 24well plates with coverslips. Wells were pre-coated with 0.2% Gelatin. Cells were allowed to settle for 48h, and then were exposed to 2h at 44°C heat stress, followed by fixation in 4% PFA. Cells were stained with ATF-6alpha (F-7) antibody (Santa Cruz, sc-166659) at 1:50 dilution, with a secondary anti-mouse antibody (Jackson Laboratories) at 1:500 dilution. Coverslips were further incubated with 1 ug/ml DAPI in PBS for 2 min before mounting. Z-stack images were taken with Zeiss LSM700 laser scanning confocal microscope. A total of 16,17,19 and 20 images were taken for Young, Young HS, Senescent and Senescent HS samples respectively, in three biological replicate experiments. An in-house MATLAB script was used in to calculate ATF6 localization, using the sum projection of z-stacks images. First, nuclei were identified in the DAPI channel by transforming values to a binary representation, and only objects bigger than a certain radius threshold were kept in order to exclude background noise. Subsequently, ATF6 channel pixels intensities were summed inside the positions previously identified as nuclei, as well as in the entire image for whole cell total intensities. Finally, for each image, the ratio of intensities was calculated (nuclei/whole cells), and data is presented in Fig. 6G, S7AI,AJ.

### Alternative splicing analysis

We used MISO (25) to examine differential alternative splicing upon HS in young and senescent cells, or between young and senescent cells. MISO GFF3 annotation files for hg19 for skipped exons, alternative 3’ and 5’ splice sites, alternative last exons, and retained introns were downloaded from the MISO website (https://miso.readthedocs.io/en/fastmiso/). GFF3 annotation file of all introns was produced using in-house scripts. All GFF3 annotations were indexed using the index_gff procedure from the MISO package. For each sample (each with two replicates), MISO was run using default settings. To assess significant differences between samples, we ran the procedure compare_miso between samples and between replicates. We denoted an event as significant if the Bayes Factors (BFs) for the comparisons between replicates were below 4 and BFs for the comparisons between different samples were greater than 8. E.g., BF of Young1 – Young2<4 and Young-HS1 – Young-HS2 < 4 and Young1 – Young-HS1>8 and Young2 – Young-HS2 >8. All reported trends were robust to different BF cutoffs, as observed in Fig. S5E-J.

### Proteasome activity assay

Young and senescent cells were plated in 6well plates and treated with HS as above. Additional recovery samples were added by moving cells from a 2h HS back to 37^0^C for 4h before lysis. Cells were washed with cold PBS and lysed in RIPA buffer (50mM Tris, 150mM NaCl, 1% NP-40, 0.5% sodium deoxycholate) with protease inhibitors and phosphatase inhibitors as above, and protein concentrations were measured using the BCA kit. Proteasome activity was quantified using the AMC-tagged proteasome fluorogenic substrate (Succ-LLVY-AMC, Calbiochem), similarly to (33). A total of 5μg protein lysate were assayed in reaction buffer (20 mM Tris-HCL (pH 7.8), 0.5 mM EDTA, 0.02% SDS) in a total volume of 100ul. MG132 was added at a final concentration of 60uM to either Young or Senescent samples, to serve as a baseline control in each experiment. Samples were incubated for 10 mins with MG132 (or DMSO) at room temp. Then, 80μM of Suc-LLVY-AMC were added and samples were transferred into 96well plates for a 30min incubation at 37^0^C. AMC fluorescence was measured using plate reader (Ex/Em 380/460 nm). For each experiment, MG132 samples fluorescence were subtracted from all other fluorescence reads (for young and senescent samples respectively), and then HS and recovery samples were divided by their respective controls.

## Acknowledgements

We thank Edith Suss-Toby and Shani Bendori from the Bioimaging center of the Faculty of Medicine Biomedical Core Facility at the Technion, for their assistance with image analysis and the Imaris software. We thank Herman Wolosker for critical reading of the manuscript. We thank the Allen and Jewel Price Center at the Technion. This project has received funding from the European Research Council under the European Union’s Horizon 2020 programme Grant 677776.

## Authors Contributions

R.S. conceived and supervised the study. F.L.A. and A.Y. performed all experiments with the help of K.R. and S.S.B. N.S. performed all computational data analyses with the help of A.M.. S.H. performed image analysis. R.S. wrote the paper with help from N.S. and input from all other authors.

## Data Availability

All RNA-seq and ribosome footprint profiling data were deposited in GEO, GSE124609.

## Supplementary Information

Supplementary Information file containing Figures S1-S9.

Supplementary Tables : Tables S1-S5.

