## Supplementary Figures for "Cellular proteostasis decline in human senescence"

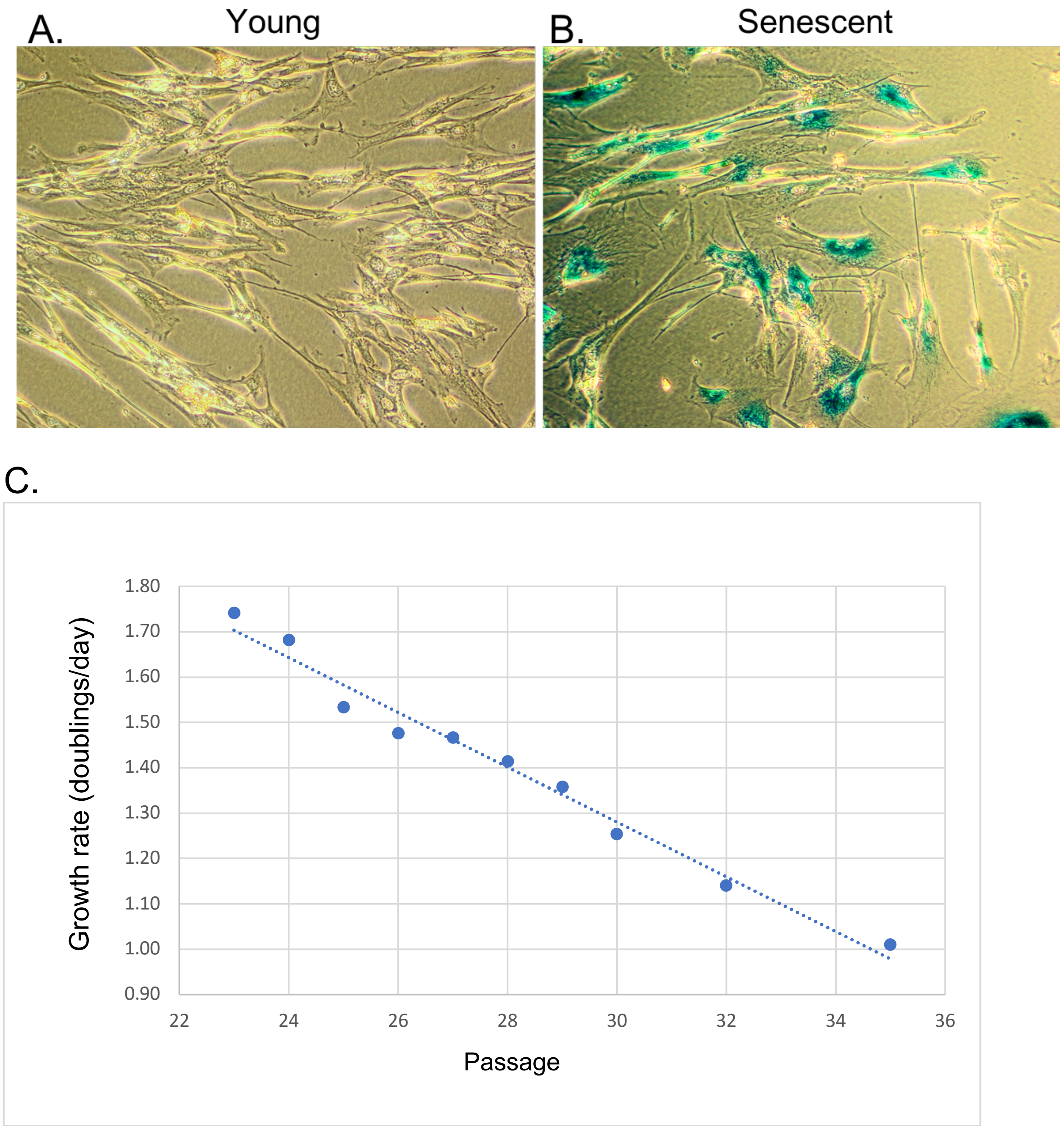


**Figure S1: Human WI38 fibroblasts grown to replicative senescence**
(A-B) SA beta-gal staining of young, passage 24 (A) vs. Senescent, p36 (B) WI38 cells demonstrate senescence. (C) Cells were counted at different passages and growth rates were. At passage 35 almost all cells have reached senescence, as seen by doubling rate of 1.01.

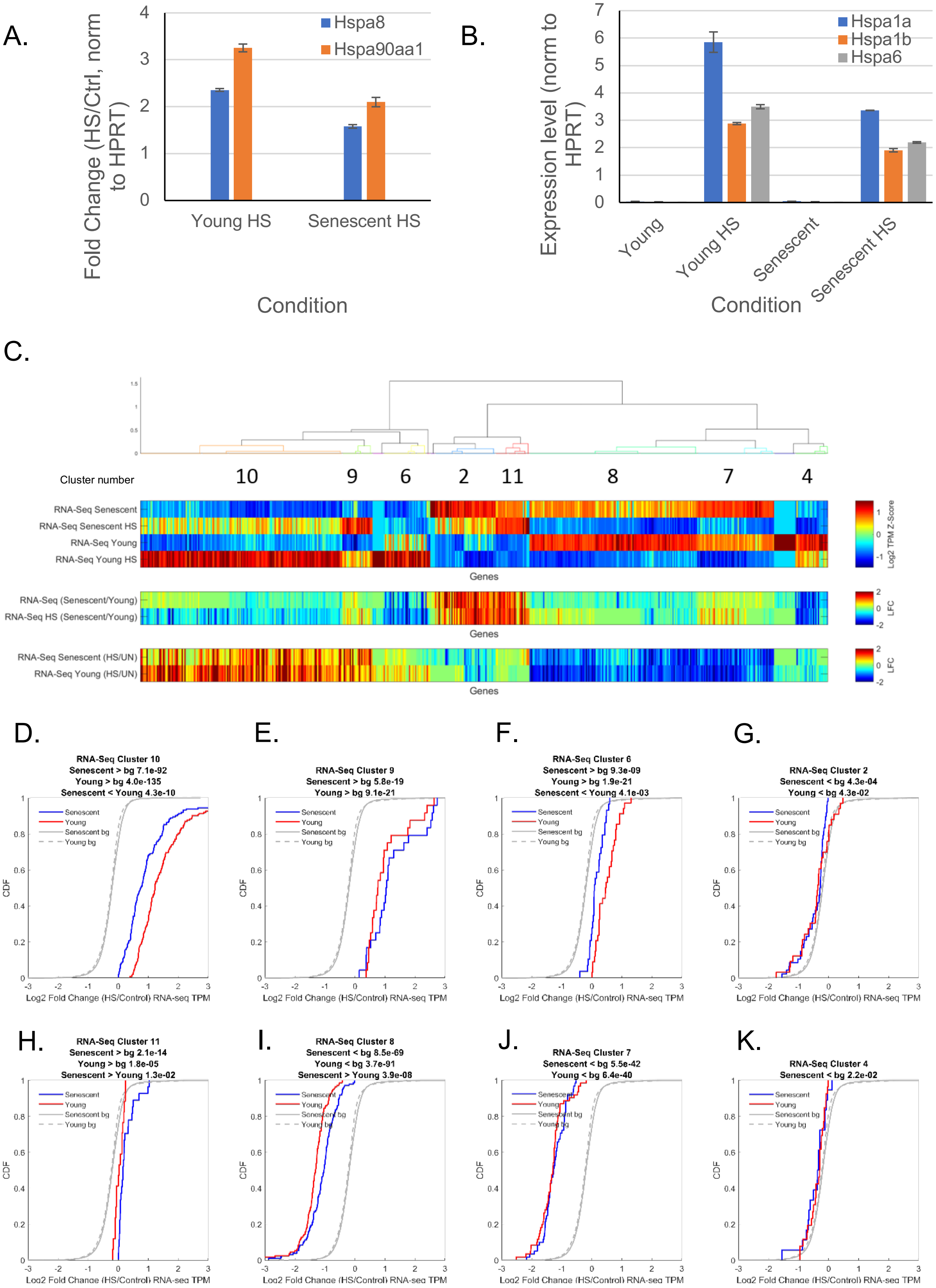
A.
Fold Change (HS/Ctrl, norm to HPRT)
Hspa8
Hspa90aa1
Young HS
Senescent HS
Condition
B.
Expression level (norm to HPRT)
Hspa1a
Hspa1b
Hspa6
Young
Young HS
Senescent
Senescent HS
Condition
C.
Cluster number
10
9
6
2
11
8
7
4
RNA-Seq Senescent
RNA-Seq Senescent HS
RNA-Seq Young
RNA-Seq Young HS
Genes
Log2 TPM Z-Score
RNA-Seq (Senescent/Young)
RNA-Seq HS (Senescent/Young)
Genes
LFC
RNA-Seq Senescent (HS/UN)
RNA-Seq Young (HS/UN)
Genes
LFC
D.
RNA-Seq Cluster 10
Senescent > bg 7.1e-92
Young > bg 4.0e-135
Senescent < Young 4.3e-10
E.
RNA-Seq Cluster 9
Senescent > bg 5.8e-19
Young > bg 9.1e-21
F.
RNA-Seq Cluster 6
Senescent > bg 9.3e-09
Young > bg 1.9e-21
Senescent < Young 4.1e-03
G.
RNA-Seq Cluster 2
Senescent < bg 4.3e-04
Young < bg 4.3e-02
H.
RNA-Seq Cluster 11
Senescent > bg 2.1e-14
Young > bg 1.8e-05
Senescent > Young 1.3e-02
I.
RNA-Seq Cluster 8
Senescent < bg 8.5e-69
Young < bg 3.7e-91
Senescent > Young 3.9e-08
J.
RNA-Seq Cluster 7
Senescent < bg 5.5e-42
Young < bg 6.4e-40
K.
RNA-Seq Cluster 4
Senescent < bg 2.2e-02
Senescent
Young
Senescent bg
Young bg
CDF
Log2 Fold Change (HS/Control) RNA-seq TPM

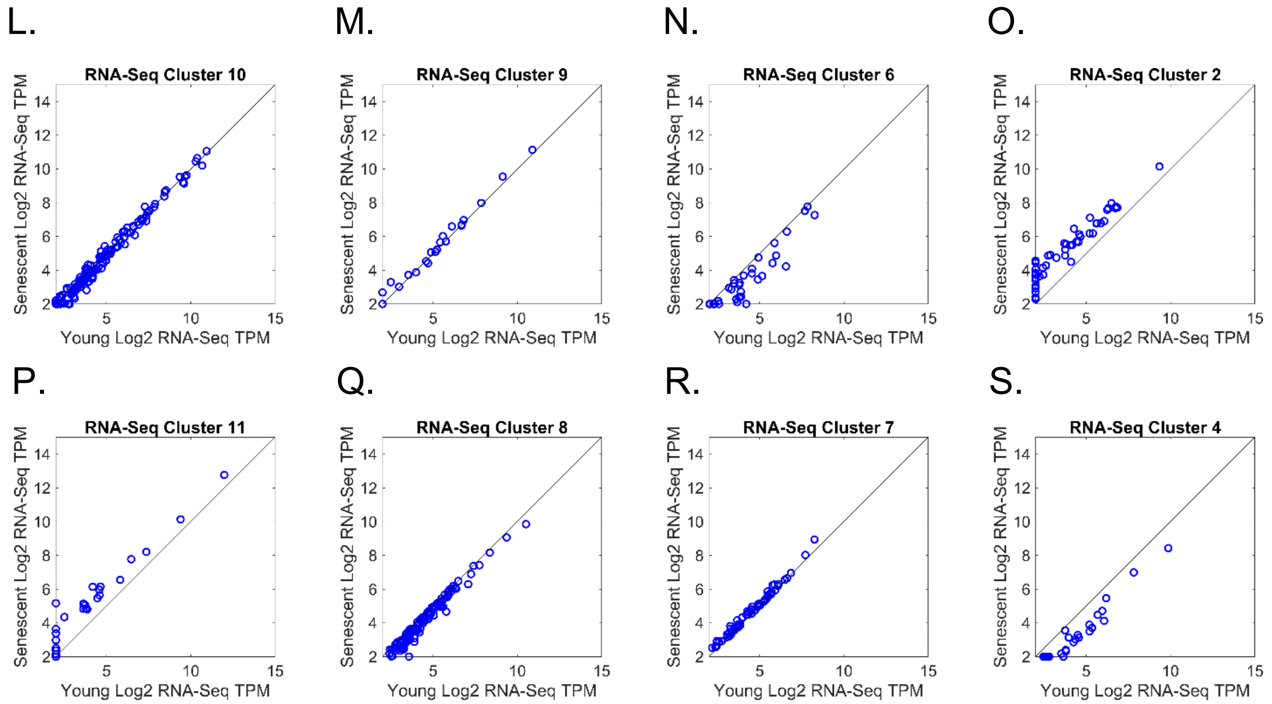


**Figure S2: Clustering analysis of differentially-expressed mRNAs in young and senescent cells subject to heat shock (HS).**

(A,B) Relative expression of representative HSP mRNAs, in young and senescent cells without or following HS were measured using realtime qPCR, represented as expression fold changes (A) for the constitutively expressed Hspa8 and Hsp90aa1, and expression levels (B) for the inducible mRNAs of Hspa1a, Hspa1b and Hspa6, whose basal expression levels before heat shock are close to zero and therefore fold changes are artificial. (C) Hierarchical clustering analysis of 548 mRNAs with significant expression difference (using DESeq2 FDR-corrected p-value < 0.05) in at least one comparison between different RNA-Seq samples. The second panel shows a gene-wise normalized Z-score heatmap of the log2 TPM scores. The upper panel shows a dendrogram resulting from the clustering procedure, performed using MATLAB. Numbers below the dendrogram note the different cluster numbers (see also Table S2), for which further information is presented below. The third and fourth panels show the log2 fold-change of senescent/young and HS/control samples, respectively. (D-K) CDF plot of the log2 expression fold change (HS/Control RNA-Seq TPM) for mRNAs in the eight major clusters within the differentially-expressed mRNAs identified by DESeq2 in the RNA-Seq exaperiments. Senescent samples - blue and young samples - red. Gray lines depict the background distributions (bg), corresponding to all expressed mRNAs in senescent (solid line) or young (dashed line) cells. p-values for significant differences between the cluster mRNAs log2 fold changes in young or senescent vs. their corresponding bg, or in young vs. senescent distributions, were calculated using KS test, and indicated above each cluster subplot. Non-significant p-values (<0.01) are not indicated. (L-S) Scatter plots of mRNA expression (log2 RNA-seq TPM) in young vs. senescent cells of the mRNAs in the eight major RNA-seq clusters.

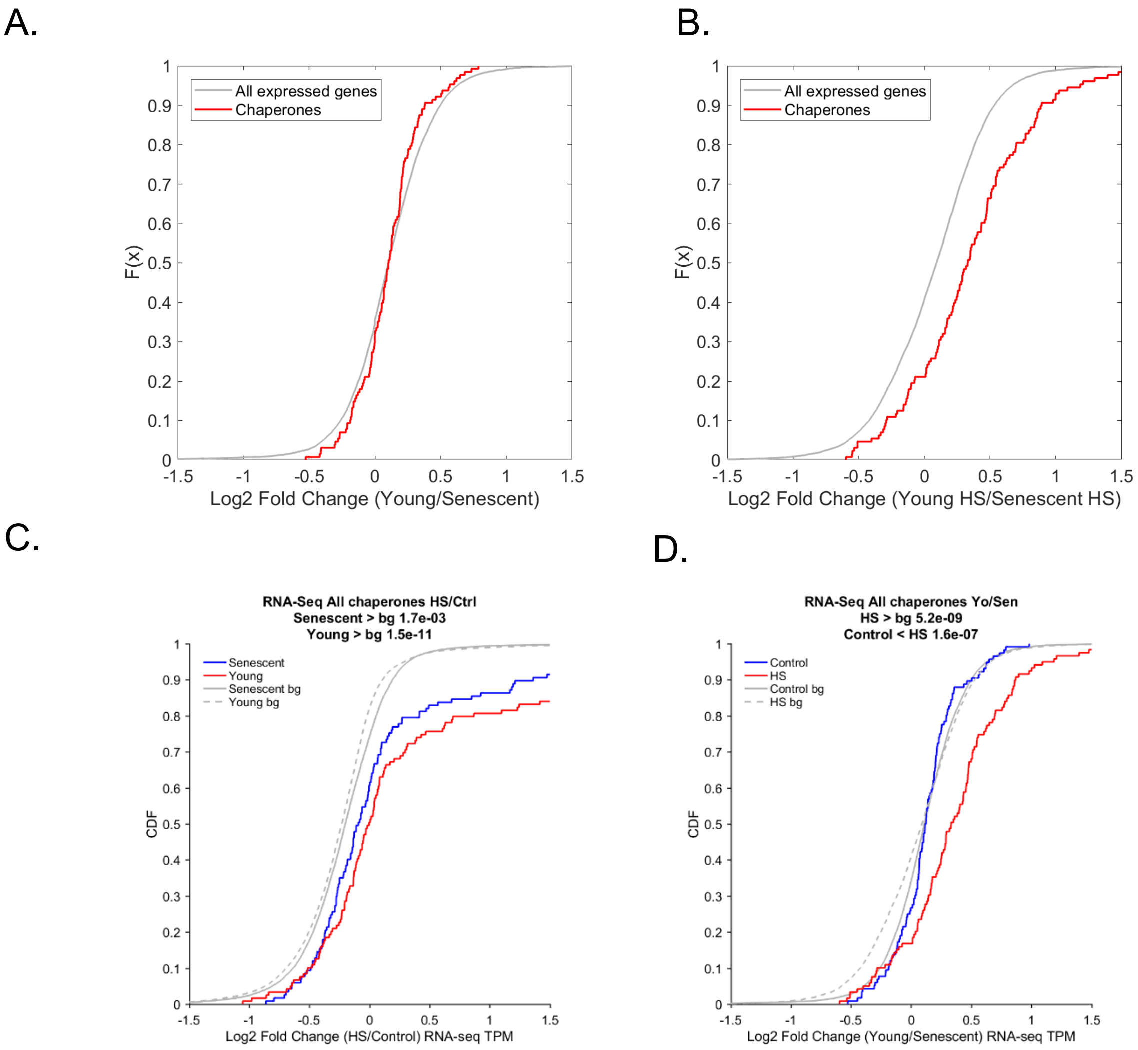
A.
B.
C.
D.
All expressed genes
Chaperones
F(x)
Log2 Fold Change (Young/Senescent)
Log2 Fold Change (Young HS/Senescent HS)
RNA-Seq All chaperones HS/Ctrl
Senescent > bg 1.7e-03
Young > bg 1.5e-11
Senescent
Young
Senescent bg
Young bg
CDF
Log2 Fold Change (HS/Control) RNA-seq TPM
RNA-Seq All chaperones Yo/Sen
HS > bg 5.2e-09
Control < HS 1.6e-07
Control
HS
Control bg
HS bg
Log2 Fold Change (Young/Senescent) RNA-seq TPM

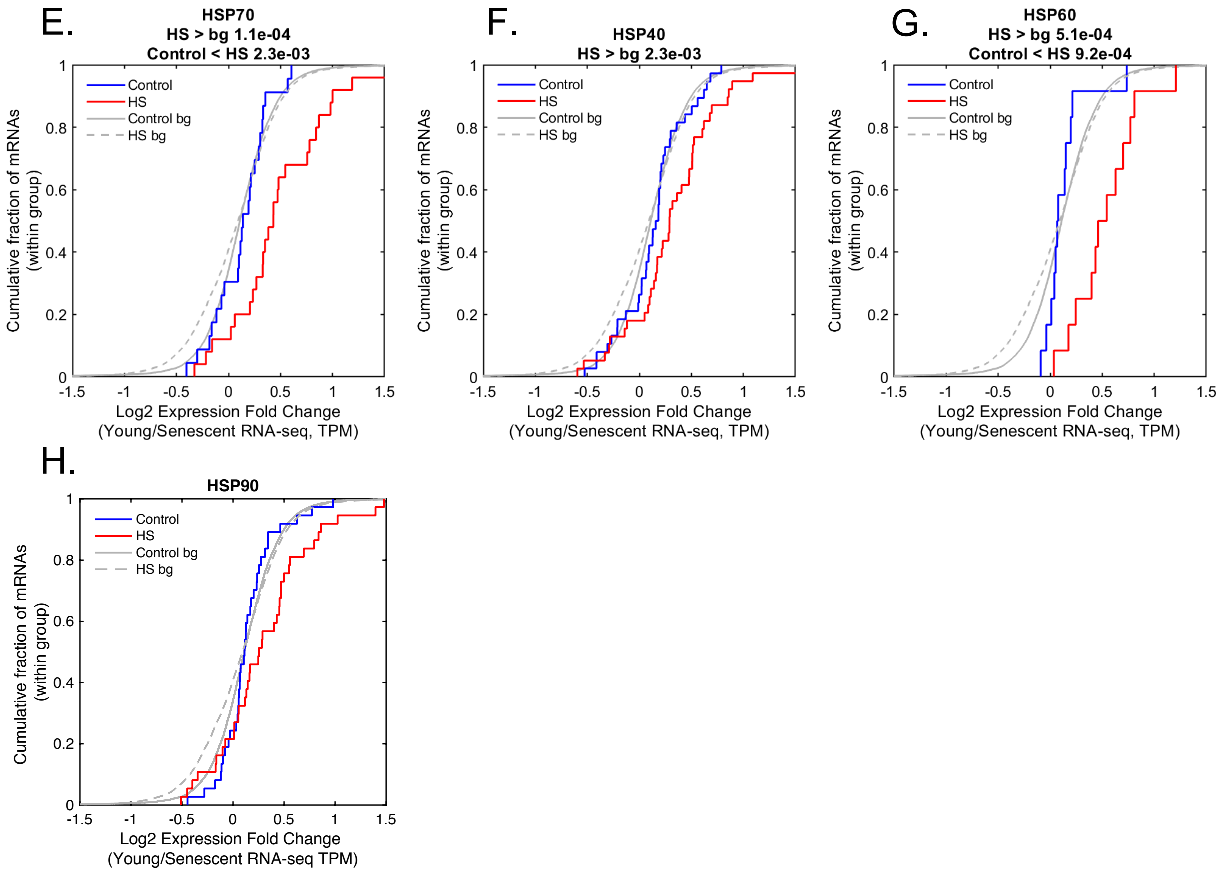


**Figure S3: Chaperone expression analysis shows impaired chaperone induction upon HS in senescent cells.**

(A-B) CDF of Log2 expression fold change (Young/Senescent, RNA-seq TPM) shows that chaperones (red) have no basal difference between unstressed young and senescent cells (A, p = 0.14, KS test), however an increased induction in young-HS relative to senescent HS is observed (B, p = $3.4^{-8}$, KS test). (C-D) CDF plots depict the log2 expression fold changes in Young/Senescent (right) or HS/Ctrl (left), for the chaperones defined in Brehme et al. (1), including all cytosolic chaperones excluding the PFD family and the TPR families, which were not changing upon HS or senescence. Cytosolic chaperones, as a group, show the proteostasis decline behavior, whereby their HS induction in senescent cells is significantly lower. P-values are indicated. (E-H) As in D, but for separate chaperone families, HSP70, HSP40, and HSP60, showing they are all demonstrating the proteostasis decline behavior. HSP90 chaperones also showed a shift, however it was not statistically significant.

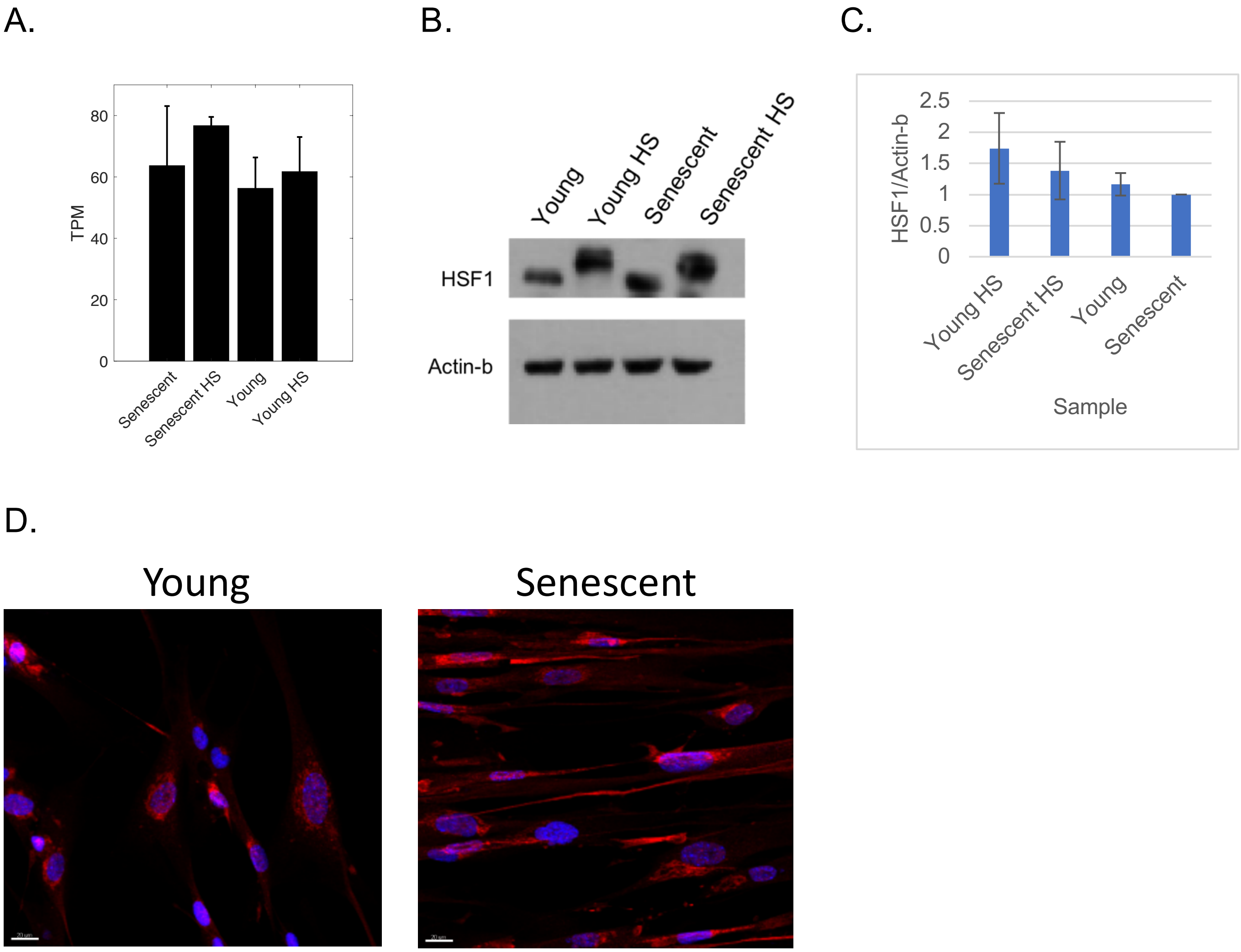
A.
80
60
40
20
0
TPM
Senescent
Senescent HS
Young
Young HS
B.
Young
Young HS
Senescent
Senescent HS
HSF1
Actin-b
C.
2.5
2
1.5
1
0.5
0
HSF1/Actin-b
Young HS
Senescent HS
Young
Senescent
Sample
D.
Young
Senescent

E.

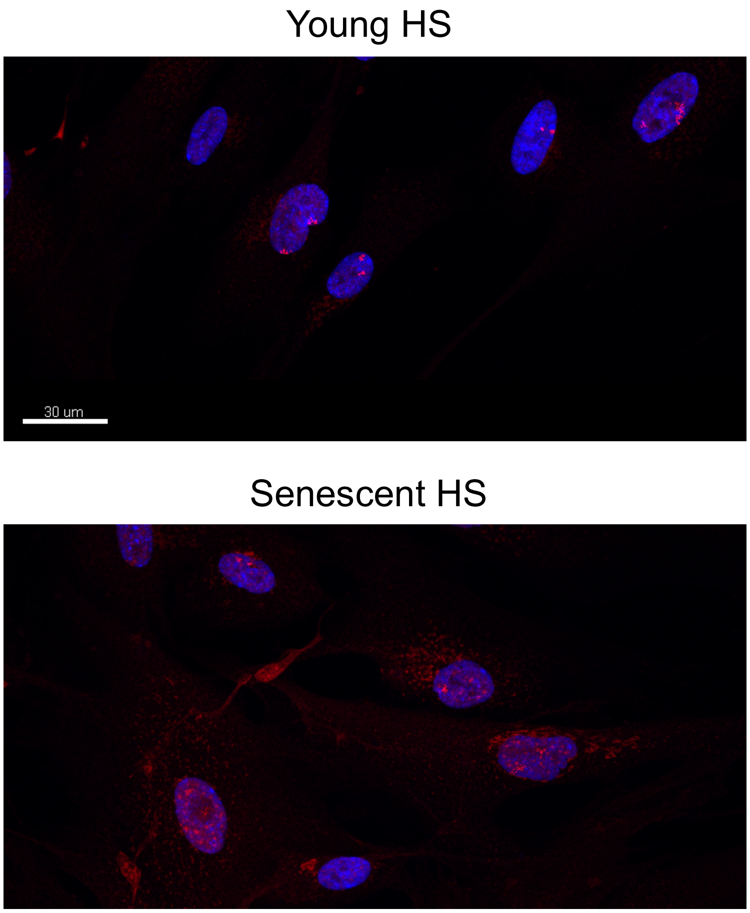


F.

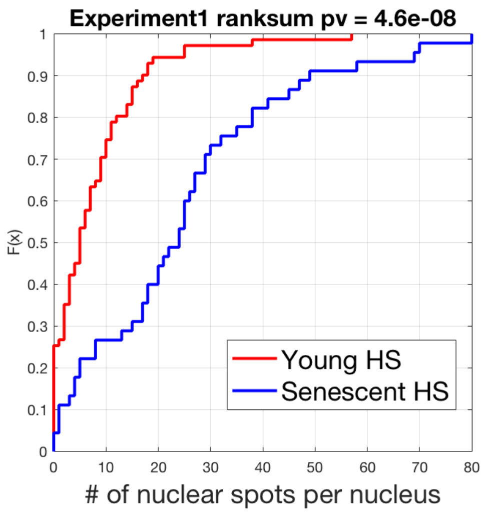


G.

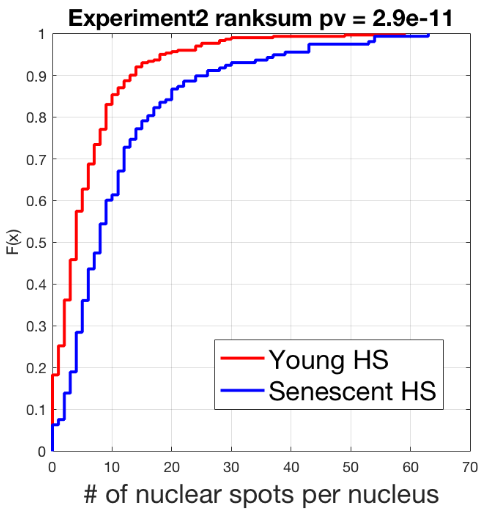


**Figure S4: phospho-HSF1 shows aberrant nuclear localization and distribution in senescent cells upon heat shock.**

(A) HSF1 mRNA levels in the different conditions. Mean of TPM expression values and std for the replicate experiments are shown, according to the RNA-seq data. (B) Biological replicate for Fig. 3A, total HSF1 levels measured using WB. (C) Quantification of the WB images of Fig. 3A and S4B were done using Fiji and HSF1 densitometry levels were normalized to the corresponding Actin-b levels. To allow comparison between experiments, all samples were normalized to the unstressed senescent sample in both experiments. Average and standard error of 4 biological replicates are presented. (D) Confocal microscopy images of unstressed young and senescent cells stained for phospho-HSF1. DAPI stain was used to mark nuclei. (E) Confocal microscopy (3D) images of phospho-HSF1 in young and senescent HS cells, as in Fig. 3C, additional fields. (F,G) CDF of the number of nuclear spots per nucleus in young (red) and senescent (blue) cells show higher number of spots per nucleus in young. (F) replicate 1, ($p = 4.6^{-8}$, ranksum test); (G) replicate 2, ($p = 2.9^{-11}$, ranksum test). Each independent replicate quantification was based on the analysis of nuclei from 6-8 fields of each of young HS and senescent HS cells.

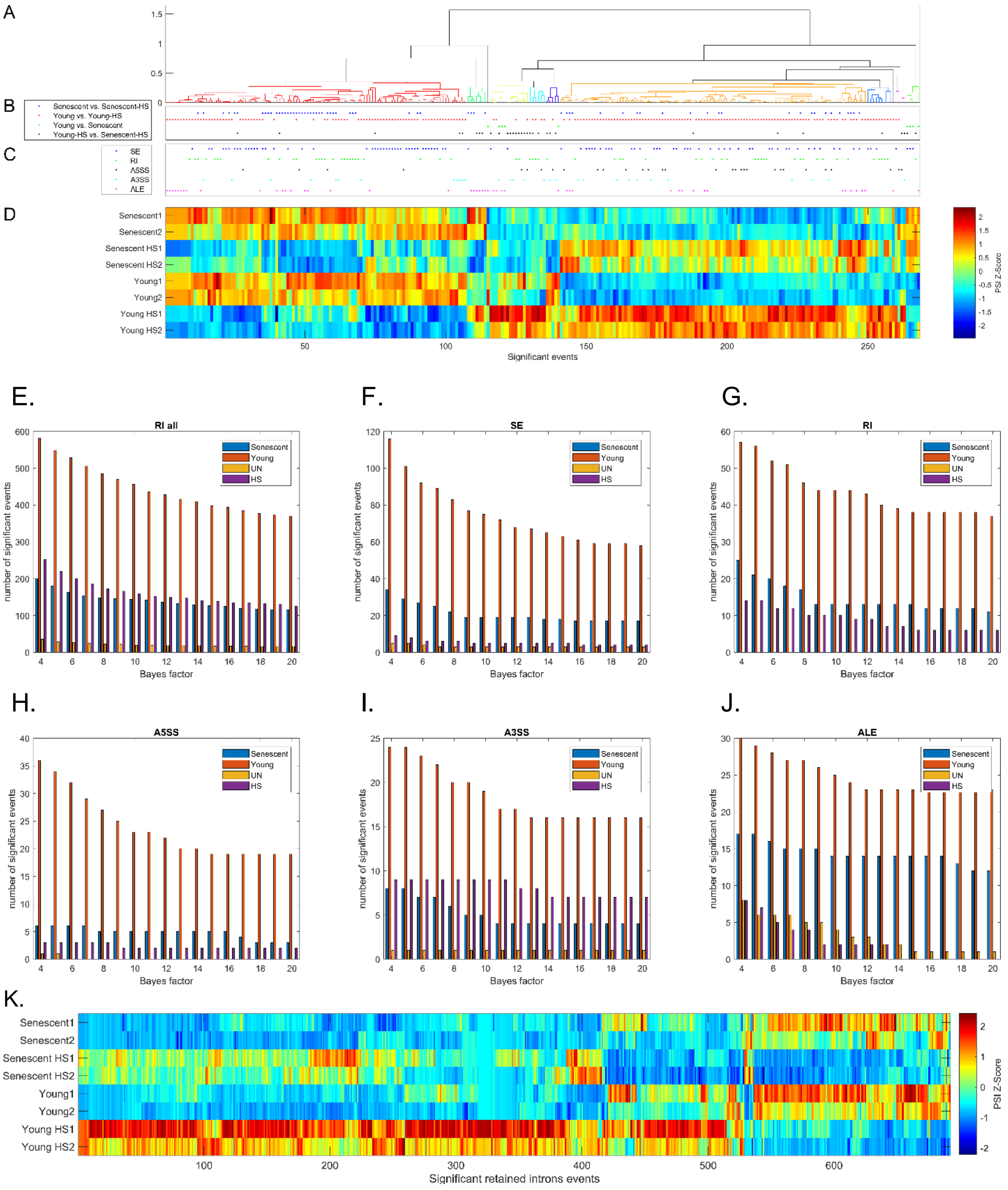


**Figure S5: Alternative splicing analyses shows deteriorated alternative splicing regulation upon HS in senescent cells.**

(A-D) Hierarchical clustering analysis of 268 significantly differential alternative splicing events in at least one comparison. (A) a dendrogram resulting from the clustering procedure. (B) Dots mark the comparison in which the event was found to be significant. (C) Dots mark the type of alternative splicing event: SE - Skipped Exon, RI - retained introns, A5SS/A3SS – alternative 5`/3` splice site, ALE – alternative last exon. (D) Heatmap of event-wise Z-score normalized percent spliced isoform (PSI) scores, calculated using MISO (2). PSI values show a robust change upon HS in young cells, while changes are much weaker in senescent cells. (E-J) The number of significant events with respect to Bayes factor (BF) cutoffs in each event-type and comparison. To assess significant differences between samples, we used the MISO algorithm (2) and ran the procedure compare_miso between samples and between replicates. We denoted an event as significant if the BFs for the comparisons between replicates were below 4 and BFs for the comparisons between samples were greater than a certain cutoff, and varied the cutoff. E.g., for a cutoff of 8 (x-axis), we plot the number of events with BF of young1 – young2 and young-HS1 – young-HS2 < 4 and young1 – young-HS1 and young2 – young-HS2 >8. The plot shows the number of significant events using a range of between-samples BFs (4-20). Blue: events of senescent vs. senescent-HS; red: events of young vs. young-HS; orange: events of young vs. senescent; purple: events of young-HS vs. senescent-HS. (K) Heatmap of event-wise Z-score normalized PSI scores resulting from hierarchical clustering analysis of 692 significantly retained introns identified in at least one comparison. PSI values show a robust change upon HS in young cells, while changes are much weaker in senescent cells.

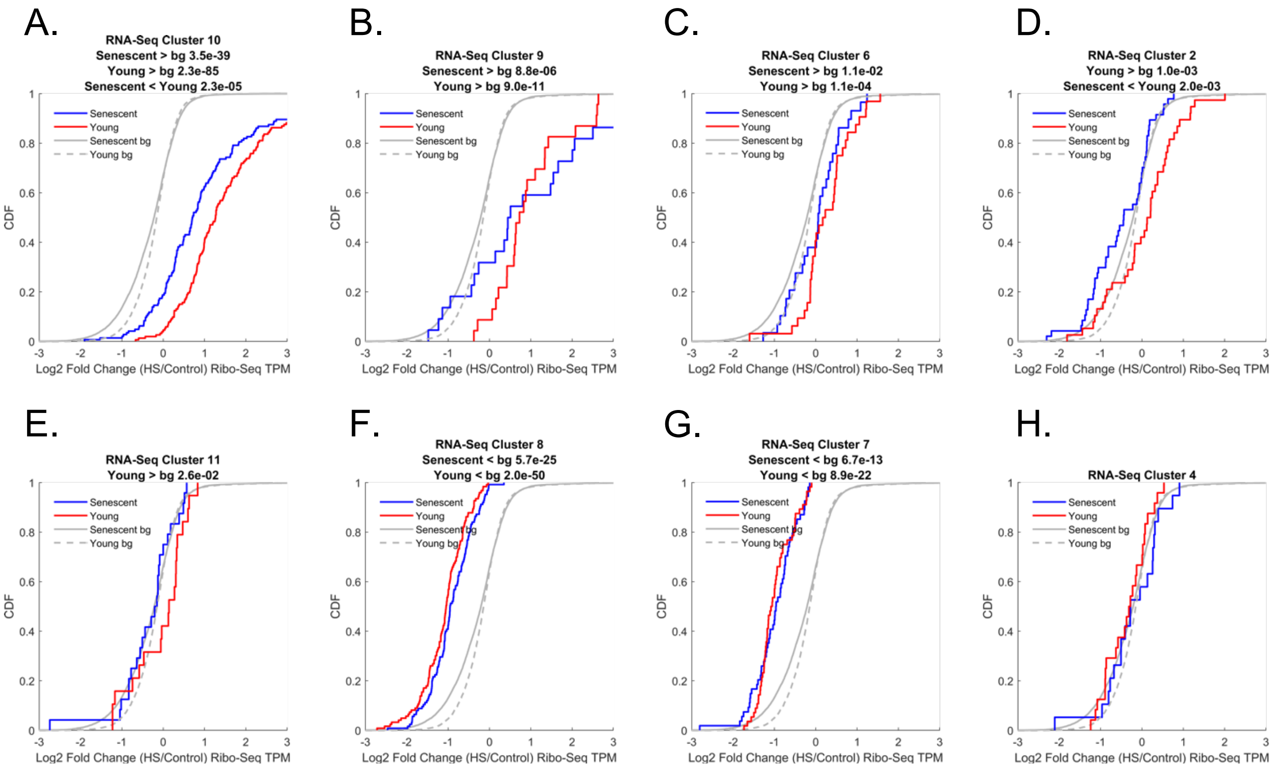
A.
RNA-Seq Cluster 10
Senescent > bg 3.5e-39
Young > bg 2.3e-85
Senescent < Young 2.3e-05
B.
RNA-Seq Cluster 9
Senescent > bg 8.8e-06
Young > bg 9.0e-11
C.
RNA-Seq Cluster 6
Senescent > bg 1.1e-02
Young > bg 1.1e-04
D.
RNA-Seq Cluster 2
Young > bg 1.0e-03
Senescent < Young 2.0e-03
E.
RNA-Seq Cluster 11
Young > bg 2.6e-02
F.
RNA-Seq Cluster 8
Senescent < bg 5.7e-25
Young < bg 2.0e-50
G.
RNA-Seq Cluster 7
Senescent < bg 6.7e-13
Young < bg 8.9e-22
H.
RNA-Seq Cluster 4
Senescent
Young
Senescent bg
Young bg
CDF
Log2 Fold Change (HS/Control) Ribo-Seq TPM

I.

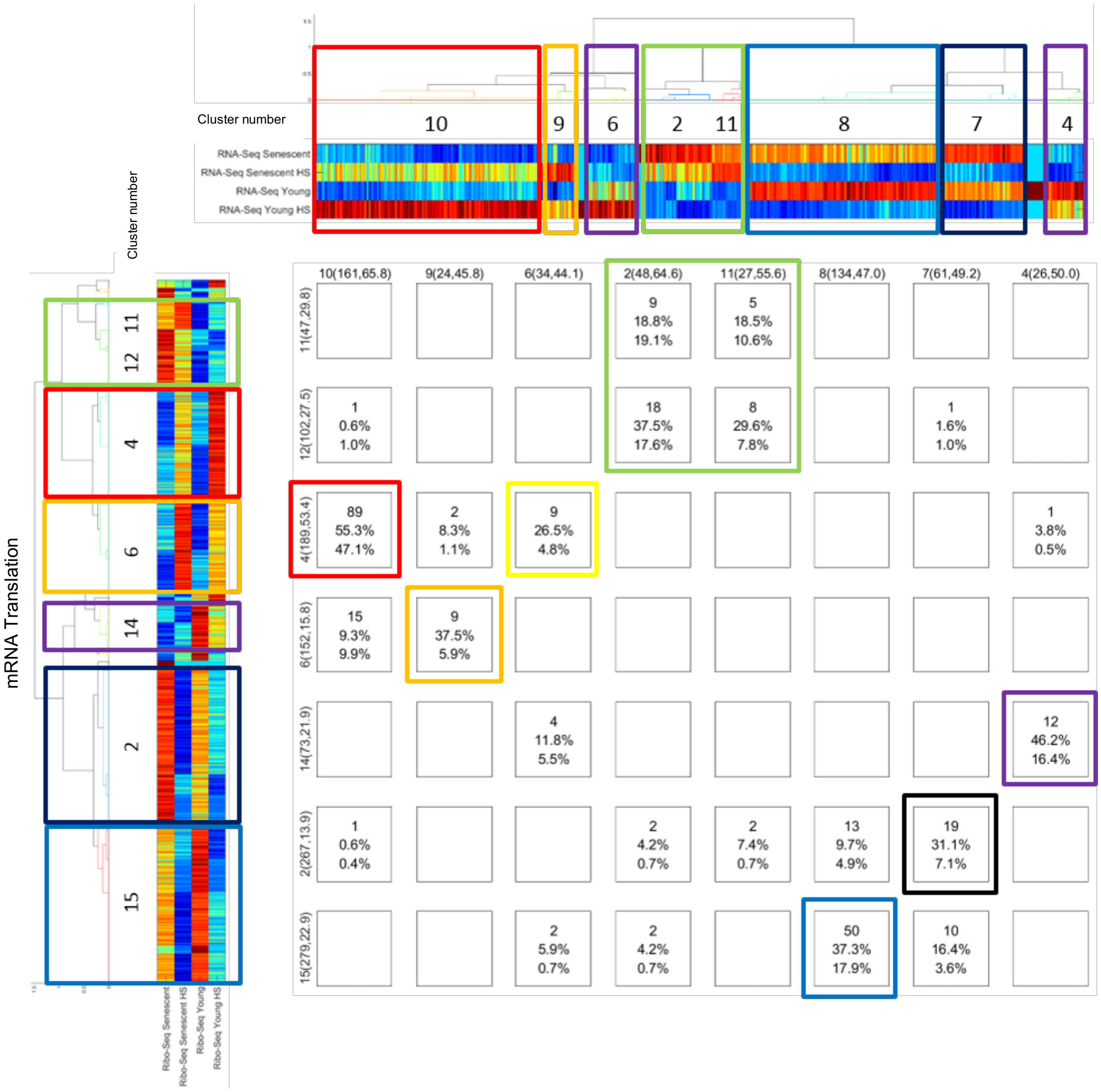


| | 10(161,65.8) | 9(24,45.8) | 6(34,44.1) | 2(48,64.6) | 11(27,55.6) | 8(134,47.0) | 7(61,49.2) | 4(26,50.0) |
|---|---|---|---|---|---|---|---|---|
| 11(47,29.8) | | | | 9<br>18.8%<br>19.1% | 5<br>18.5%<br>10.6% | | | |
| 12(102,27.5) | 1<br>0.6%<br>1.0% | | | 18<br>37.5%<br>17.6% | 8<br>29.6%<br>7.8% | | 1<br>1.6%<br>1.0% | |
| 4(189,53.4) | 89<br>55.3%<br>47.1% | 2<br>8.3%<br>1.1% | 9<br>26.5%<br>4.8% | | | | | 1<br>3.8%<br>0.5% |
| 6(152,15.8) | 15<br>9.3%<br>9.9% | 9<br>37.5%<br>5.9% | | | | | | |
| 14(73,21.9) | | | 4<br>11.8%<br>5.5% | | | | | 12<br>46.2%<br>16.4% |
| 2(267,13.9) | 1<br>0.6%<br>0.4% | | | 2<br>4.2%<br>0.7% | 2<br>7.4%<br>0.7% | 13<br>9.7%<br>4.9% | 19<br>31.1%<br>7.1% | |
| 15(279,22.9) | | | 2<br>5.9%<br>0.7% | 2<br>4.2%<br>0.7% | | 50<br>37.3%<br>17.9% | 10<br>16.4%<br>3.6% | |

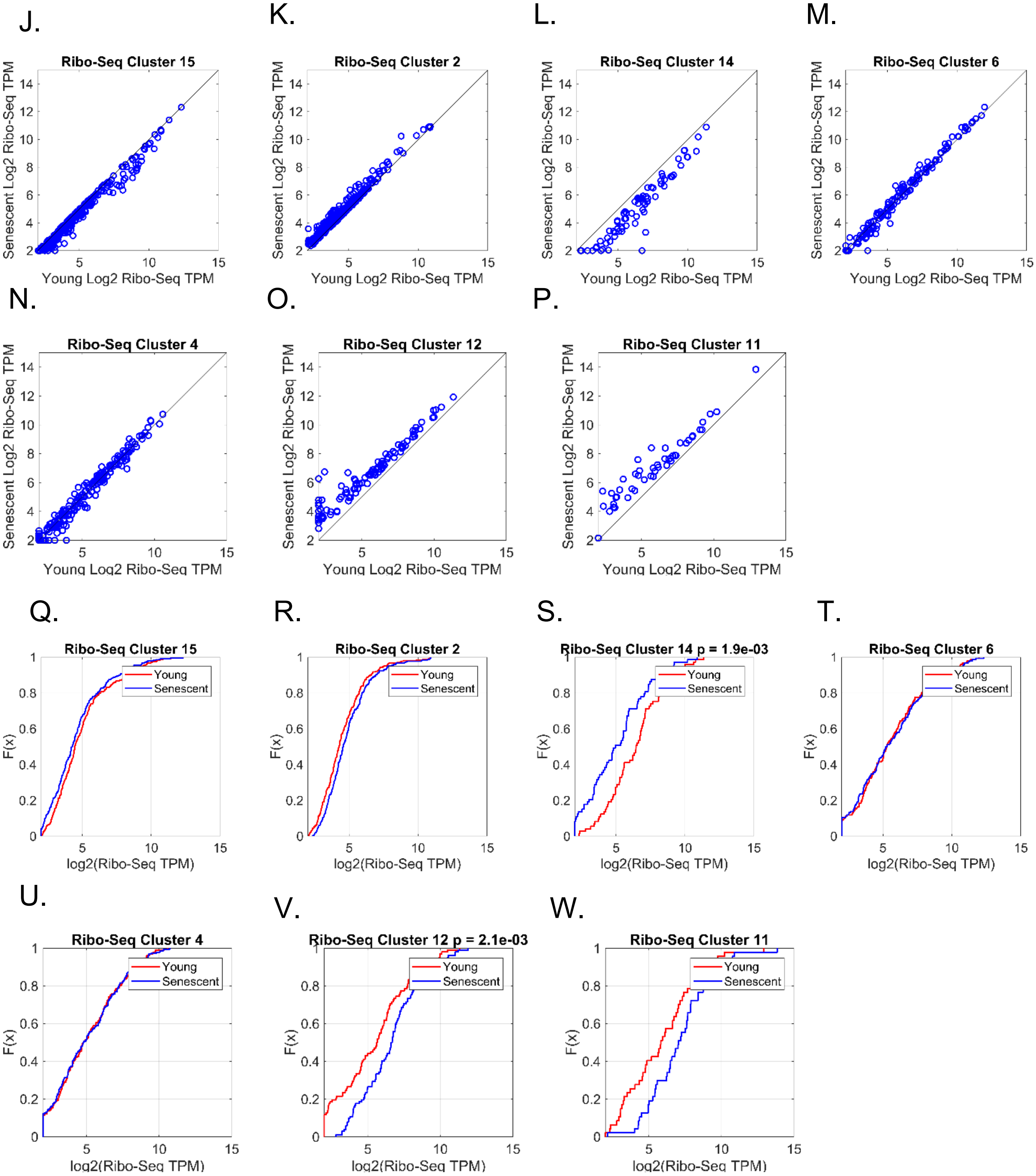

J.
Ribo-Seq Cluster 15
Senescent Log2 Ribo-Seq TPM
Young Log2 Ribo-Seq TPM
K.
Ribo-Seq Cluster 2
Senescent Log2 Ribo-Seq TPM
Young Log2 Ribo-Seq TPM
L.
Ribo-Seq Cluster 14
Senescent Log2 Ribo-Seq TPM
Young Log2 Ribo-Seq TPM
M.
Ribo-Seq Cluster 6
Senescent Log2 Ribo-Seq TPM
Young Log2 Ribo-Seq TPM
N.
Ribo-Seq Cluster 4
Senescent Log2 Ribo-Seq TPM
Young Log2 Ribo-Seq TPM
O.
Ribo-Seq Cluster 12
Senescent Log2 Ribo-Seq TPM
Young Log2 Ribo-Seq TPM
P.
Ribo-Seq Cluster 11
Senescent Log2 Ribo-Seq TPM
Young Log2 Ribo-Seq TPM
Q.
Ribo-Seq Cluster 15
Young
Senescent
F(x)
log2(Ribo-Seq TPM)
R.
Ribo-Seq Cluster 2
Young
Senescent
F(x)
log2(Ribo-Seq TPM)
S.
Ribo-Seq Cluster 14 p = 1.9e-03
Young
Senescent
F(x)
log2(Ribo-Seq TPM)
T.
Ribo-Seq Cluster 6
Young
Senescent
F(x)
log2(Ribo-Seq TPM)
U.
Ribo-Seq Cluster 4
Young
Senescent
F(x)
log2(Ribo-Seq TPM)
V.
Ribo-Seq Cluster 12 p = 2.1e-03
Young
Senescent
F(x)
log2(Ribo-Seq TPM)
W.
Ribo-Seq Cluster 11
Young
Senescent
F(x)
log2(Ribo-Seq TPM)

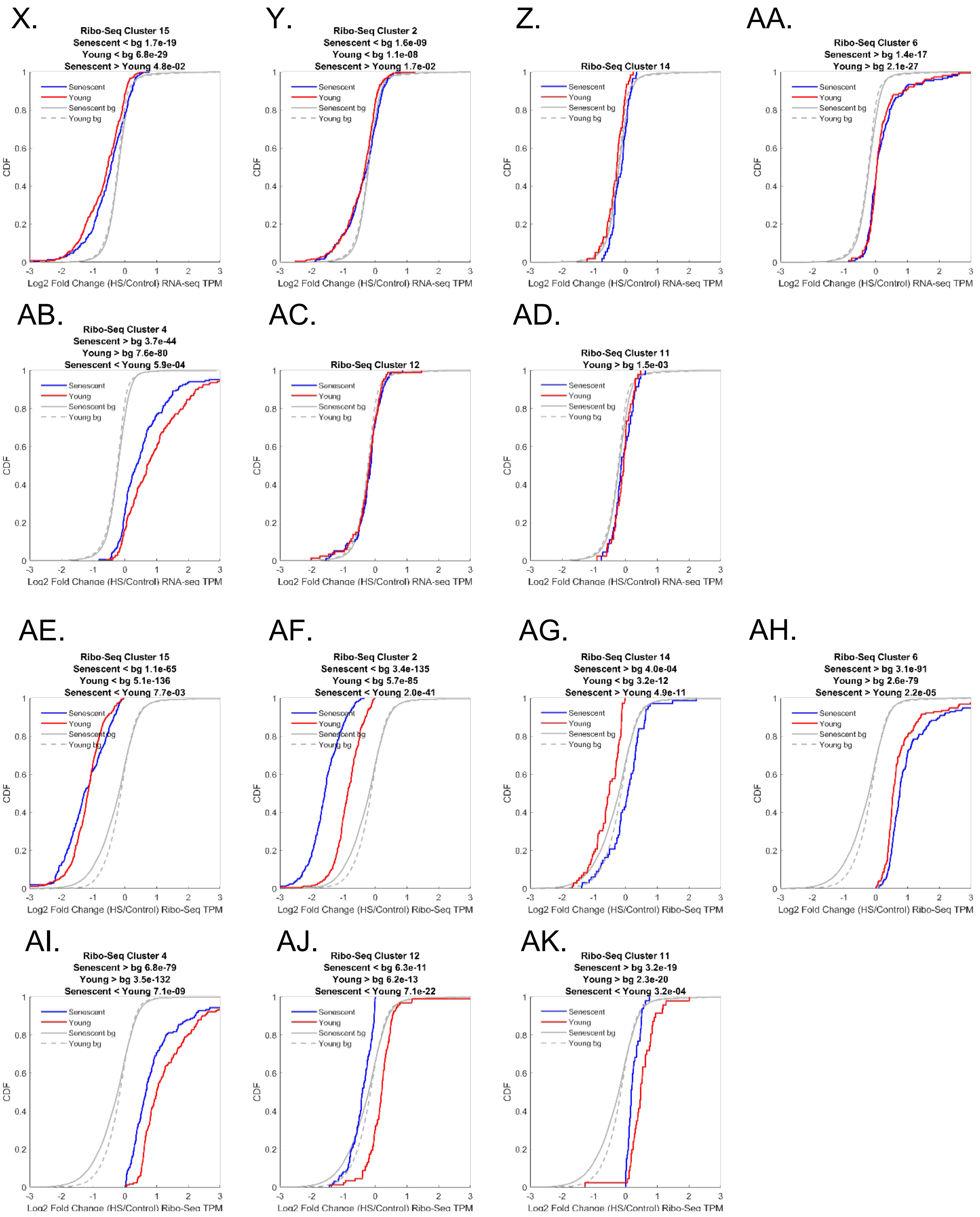


**Figure S6: Expression patterns in RNA-Seq and Ribo-Seq major clusters.** (A-H) CDF plots of the log2 translation fold change (HS/Control Ribo-Seq TPM) for mRNAs in the eight major clusters within the mRNA expression differentially-expressed mRNAs (derived from RNA-Seq, defined in Fig. S2A, Table S2A). (A-H,Q-AK) Senescent samples are marked in blue and young samples are marked in red. Gray lines depict the background distributions (bg), corresponding to all expressed mRNAs in senescent (solid line) and young (dashed line) cells. (I) Overlaps between mRNA expression (RNA-Seq, top) and translation (Ribo-Seq, left) major clusters (see Table S2A,B for cluster numbers, as also indicated here). Each cluster number is followed by its size (number of mRNAs) and the percentage of mRNAs in the intersection with all reciprocal clusters together. Inside each square we note (from top to bottom) the number of intersecting mRNAs, the percentage out of the expression (RNA-Seq) cluster, and the percentage out of the translation (Ribo-Seq) cluster. Except for translation (Ribo-Seq) cluster 4 that overlaps with expression cluster 10 (the proteostasis decline clusters, marked in red), there is little overlap of translation (Ribo-Seq) clusters with expression (RNA-Seq) clusters (below 20%), whereas RNA-Seq clusters have somewhat larger coverage within Ribo-Seq clusters (up to 46%). Clusters with matching behavior are colored by the same color. The yellow-marked square denotes the high intersection (26.5%) of expression (RNA-Seq) cluster 6 (young-specific with HS induction) with translation (Ribo-Seq) cluster 4 (HS induction, higher in young), which have different behaviors, but the overlap is very small (only 9 genes) (J-P) Scatter plots of translation levels (Ribo-Seq, log2 TPM) in unstressed young vs. senescent samples of the mRNAs in the seven major translation clusters. (Q-W) CDF plots of the log2 translation levels (Ribo-Seq TPM) in young (red) and senescent (blue) samples in the seven major translation clusters. (X-AD) CDF plots of the log2 mRNA expression fold-change (HS/Control, RNA-Seq TPM) for genes in the seven major translation clusters. (AE-AK) CDF plots of the log2 translation fold-change (HS/Control, Ribo-Seq TPM) for mRNAs in the seven major translation clusters.

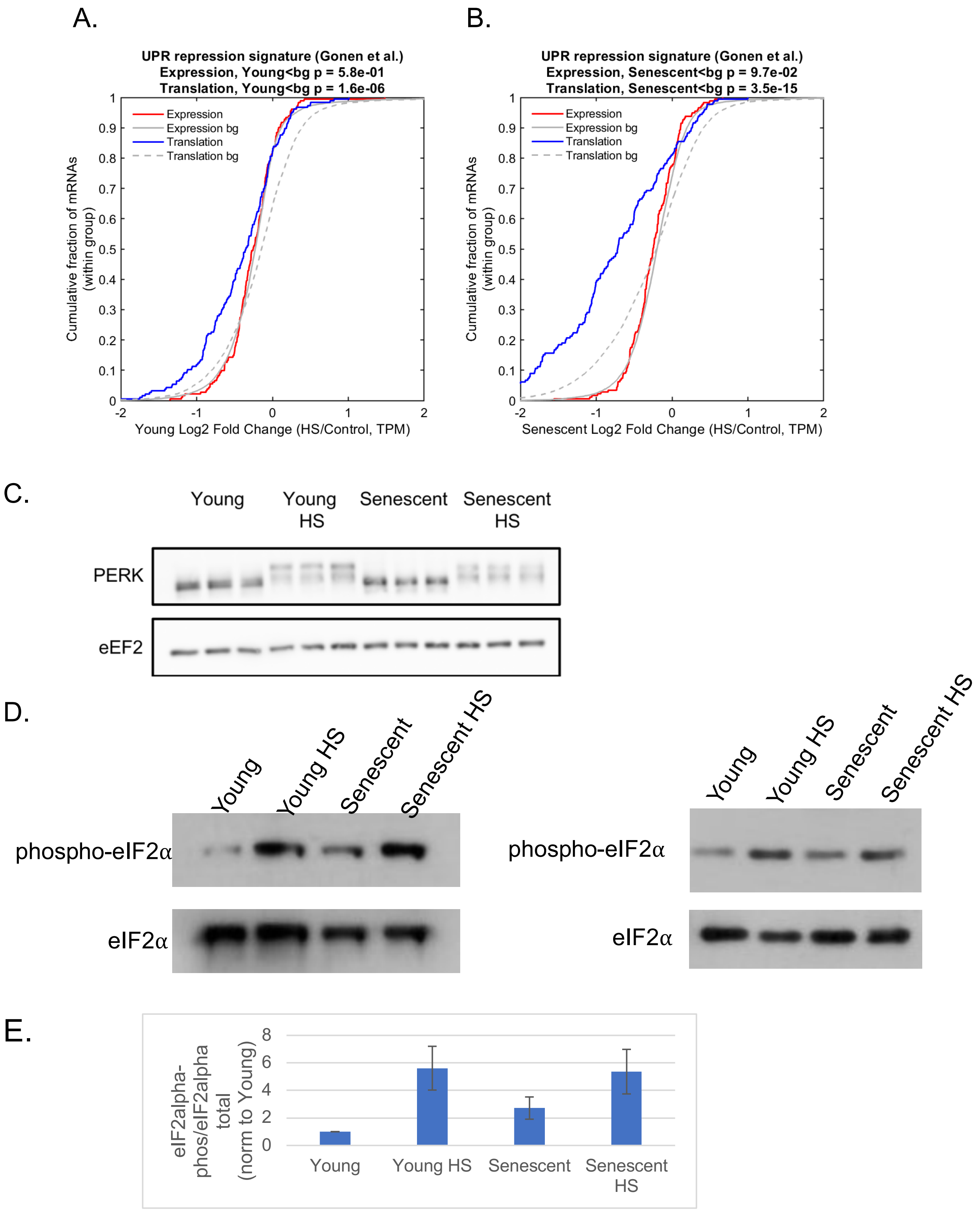
A.
UPR repression signature (Gonen et al.)
Expression, Young<bg p = 5.8e-01
Translation, Young<bg p = 1.6e-06
Expression
Expression bg
Translation
Translation bg
Cumulative fraction of mRNAs (within group)
Young Log2 Fold Change (HS/Control, TPM)
B.
UPR repression signature (Gonen et al.)
Expression, Senescent<bg p = 9.7e-02
Translation, Senescent<bg p = 3.5e-15
Senescent Log2 Fold Change (HS/Control, TPM)
C.
Young
Young HS
Senescent
Senescent HS
PERK
eEF2
D.
phospho-eIF2α
eIF2α
E.
eIF2alpha-phos/eIF2alpha total (norm to Young)
Young
Young HS
Senescent
Senescent HS

F.

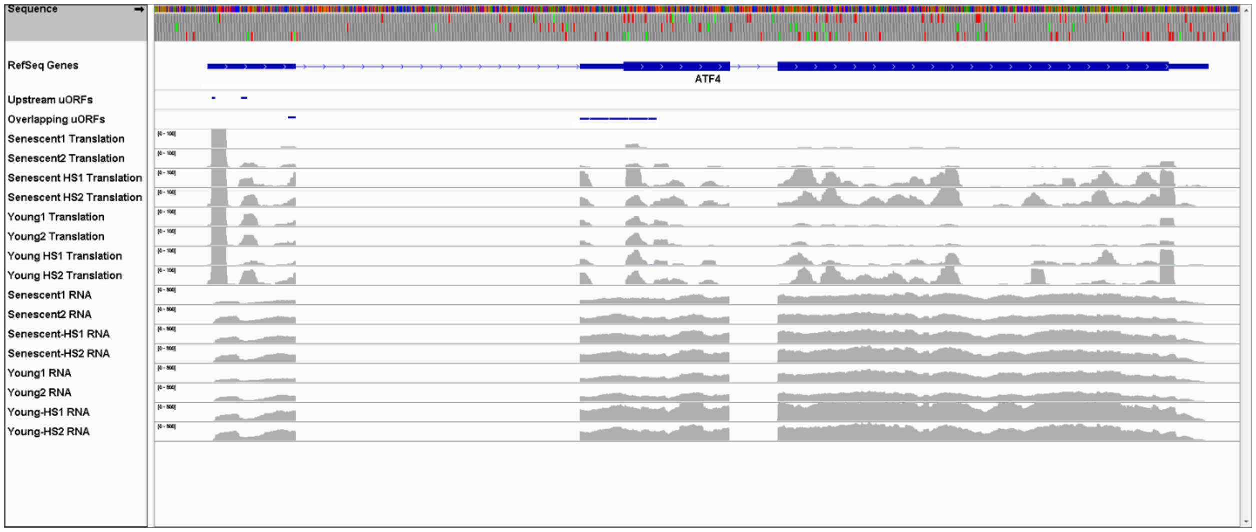
Sequence
RefSeq Genes
ATF4
Upstream uORFs
Overlapping uORFs
Senescent1 Translation
Senescent2 Translation
Senescent HS1 Translation
Senescent HS2 Translation
Young1 Translation
Young2 Translation
Young HS1 Translation
Young HS2 Translation
Senescent1 RNA
Senescent2 RNA
Senescent-HS1 RNA
Senescent-HS2 RNA
Young1 RNA
Young2 RNA
Young-HS1 RNA
Young-HS2 RNA

G.

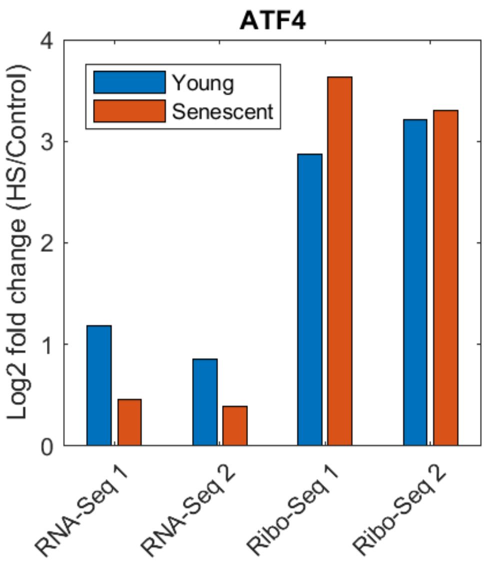
ATF4
Young
Senescent
Log2 fold change (HS/Control)
0
1
2
3
4
RNA-Seq 1
RNA-Seq 2
Ribo-Seq 1
Ribo-Seq 2

H.

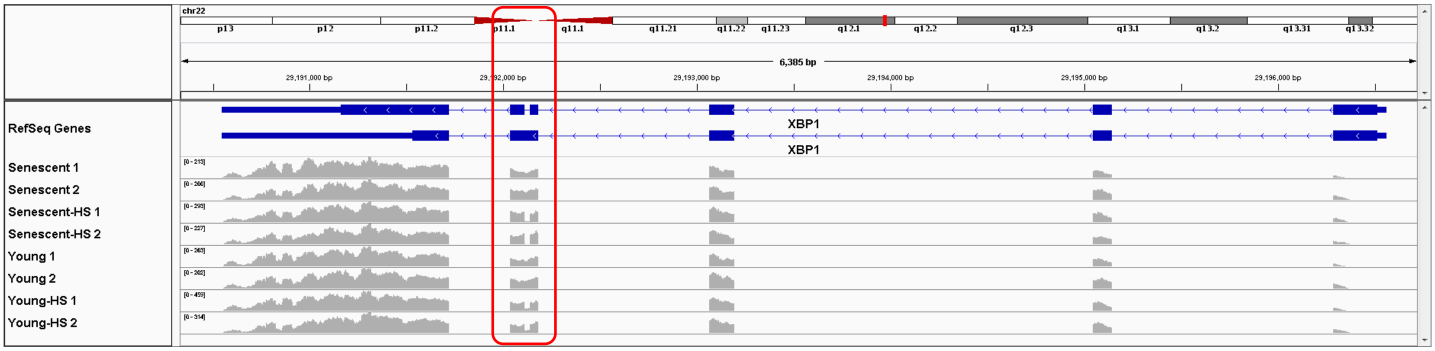
chr22
p13
p12
p11.2
p11.1
q11.1
q11.21
q11.22
q11.23
q12.1
q12.2
q12.3
q13.1
q13.2
q13.31
q13.32
6,385 bp
29,191,000 bp
29,193,000 bp
29,194,000 bp
29,195,000 bp
29,196,000 bp
RefSeq Genes
XBP1
XBP1
Senescent 1
Senescent 2
Senescent-HS 1
Senescent-HS 2
Young 1
Young 2
Young-HS 1
Young-HS 2

I.

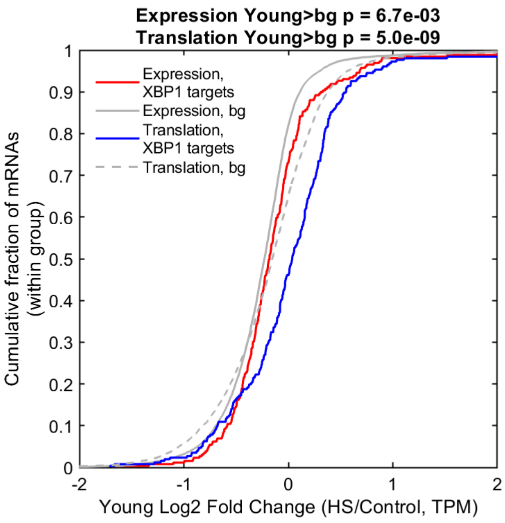
Expression Young>bg p = 6.7e-03
Translation Young>bg p = 5.0e-09
Expression, XBP1 targets
Expression, bg
Translation, XBP1 targets
Translation, bg
Cumulative fraction of mRNAs (within group)
Young Log2 Fold Change (HS/Control, TPM)

J.

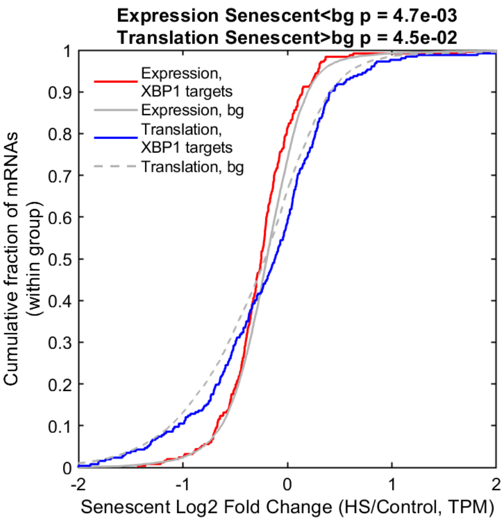
Expression Senescent<bg p = 4.7e-03
Translation Senescent>bg p = 4.5e-02
Expression, XBP1 targets
Expression, bg
Translation, XBP1 targets
Translation, bg
Cumulative fraction of mRNAs (within group)
Senescent Log2 Fold Change (HS/Control, TPM)

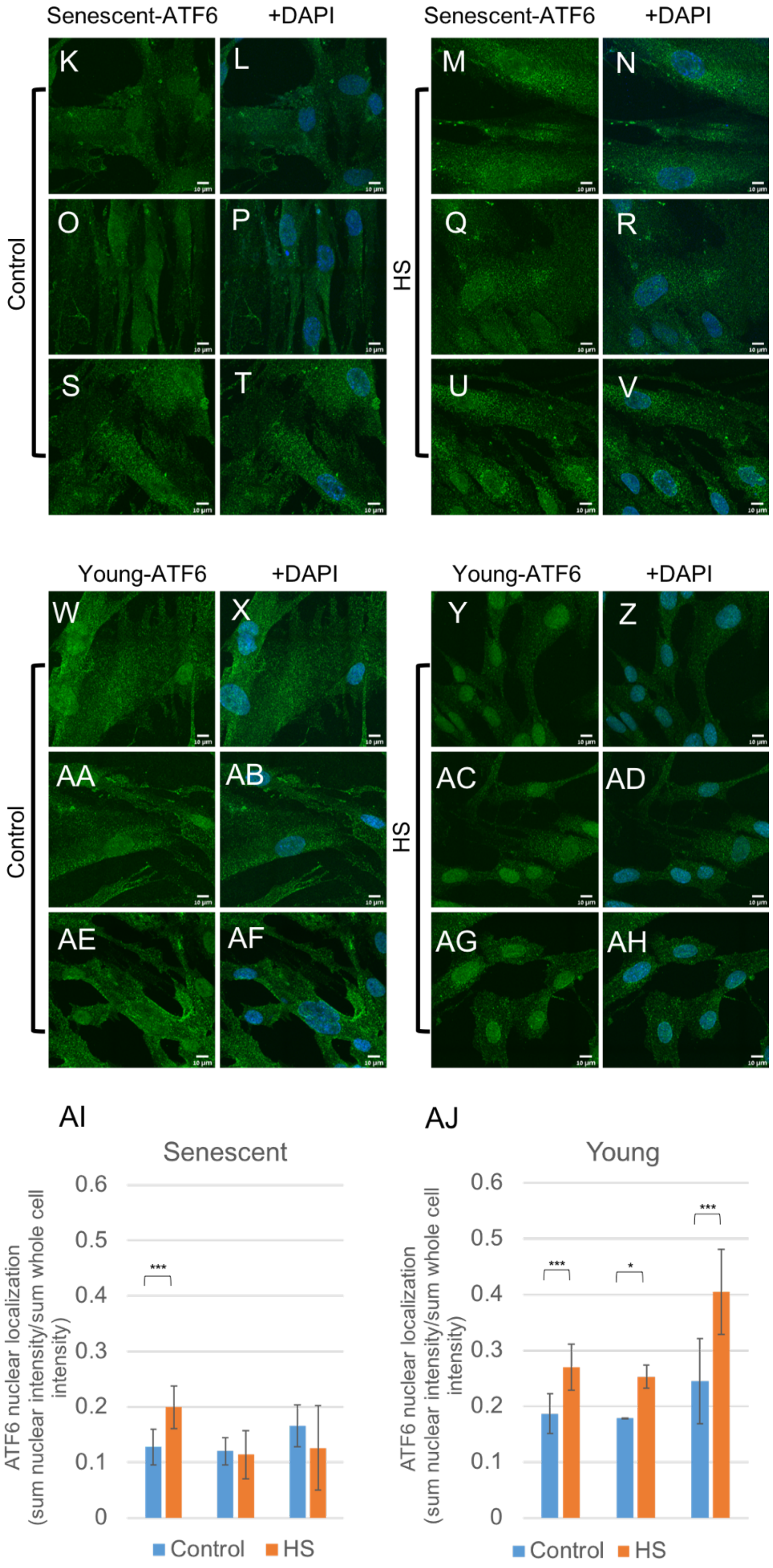
Senescent-ATF6
+DAPI
Senescent-ATF6
+DAPI
K
L
M
N
O
P
Q
R
S
T
U
V
Control
HS
Young-ATF6
+DAPI
Young-ATF6
+DAPI
W
X
Y
Z
AA
AB
AC
AD
AE
AF
AG
AH
Control
HS
AI
Senescent
ATF6 nuclear localization (sum nuclear intensity/sum whole cell intensity)
0.6
0.5
0.4
0.3
0.2
0.1
0
***
Control
HS
AJ
Young
ATF6 nuclear localization (sum nuclear intensity/sum whole cell intensity)
0.6
0.5
0.4
0.3
0.2
0.1
0
***
*
***
Control
HS

**Figure S7: Loss of coordination between UPR branches in stressed senescent cells.**
(A, B) CDF plot of the log2 fold-changes in expression (red) or translation (blue) (HS/Control, TPM) of the group of mRNAs within the UPR repression signature previously identified in Gonen et al. (3) in young (A) and senescent (B) cells. Expression (RNA-Seq, red) and translation (Ribo-Seq, blue) log2 fold change distributions are compared to background distribution of all expressed genes (grey solid and dashed lines, respectively) and demonstrate stronger repression in senescent cells at the level of translation. KS test p-values for the comparisons are indicated. (C) Western blot for PERK shows an upshift typical of phosphorylation in both young and senescent HS cells. (D) eIF2α phosphorylation (Ser51) was assayed using WB in young and senescent cells, before and after HS, as well as total eIF2α levels. Two representative experiments are shown. (E) WB signal quantification was performed using Fiji, and phospho-eIF2α levels were normalized to the levels of total eIF2α in each experiment. Then, this ratio was further normalized to the level in Young unstressed cells to allow comparison between different experiments. The graph shows the average and standard error of 4 biological replicates. (F) Visualization of the read density of translation (Ribo-Seq) samples (top) and mRNA expression (RNA-seq) samples (bottom) across the ATF4 gene locus, using the IGV tool (4). The plot demonstrates increased translation of ATF4 main ORF (RefSeq annotation presented) upon heat shock. uORF (upstream ORFs) denoted in the top annotation tracks. An orange arrow marks a low mappability region for short reads (approximate, from the hg19 wgEncodeDukeMapabilityUniqueness35bp ucsc track), explaining why no Ribo-seq reads have been mapped to the genome in that region. (G) Quantification of the induction levels of ATF4 main ORF translation level (Ribo-seq, log2 fold change HS/control), and mRNA level (RNA-seq, log2 fold change HS/control) (H) Visualization of the read density of mRNA expression (RNA-Seq) samples across the entire XBP1 gene locus. The red box marks the exon that is affected by IRE1-mediated splicing. (I,J) CDF plots of the log2 fold change (HS/Control, TPM) in mRNA expression or translation, of XBP1-s target genes (gene list taken from Shoulders et al.(5)) demonstrate their significant induction in young cells (I) and lack of significant induction in senescent cells (J). KS test p-values for the significance of induction are indicated above. (K-AH) ATF6 immunofluorescence staining, as in Fig. 6F, additional fields. (AI-AJ) As in Fig. 6G, mean and std of ATF6 nuclear localization, calculated as the sum of fluorescence intensity in the nuclei divided by the sum of fluorescence intensity in entire cells (see Methods), for the three biological replicates separately. ***: $p<0.005$, *:$0<0.05$.

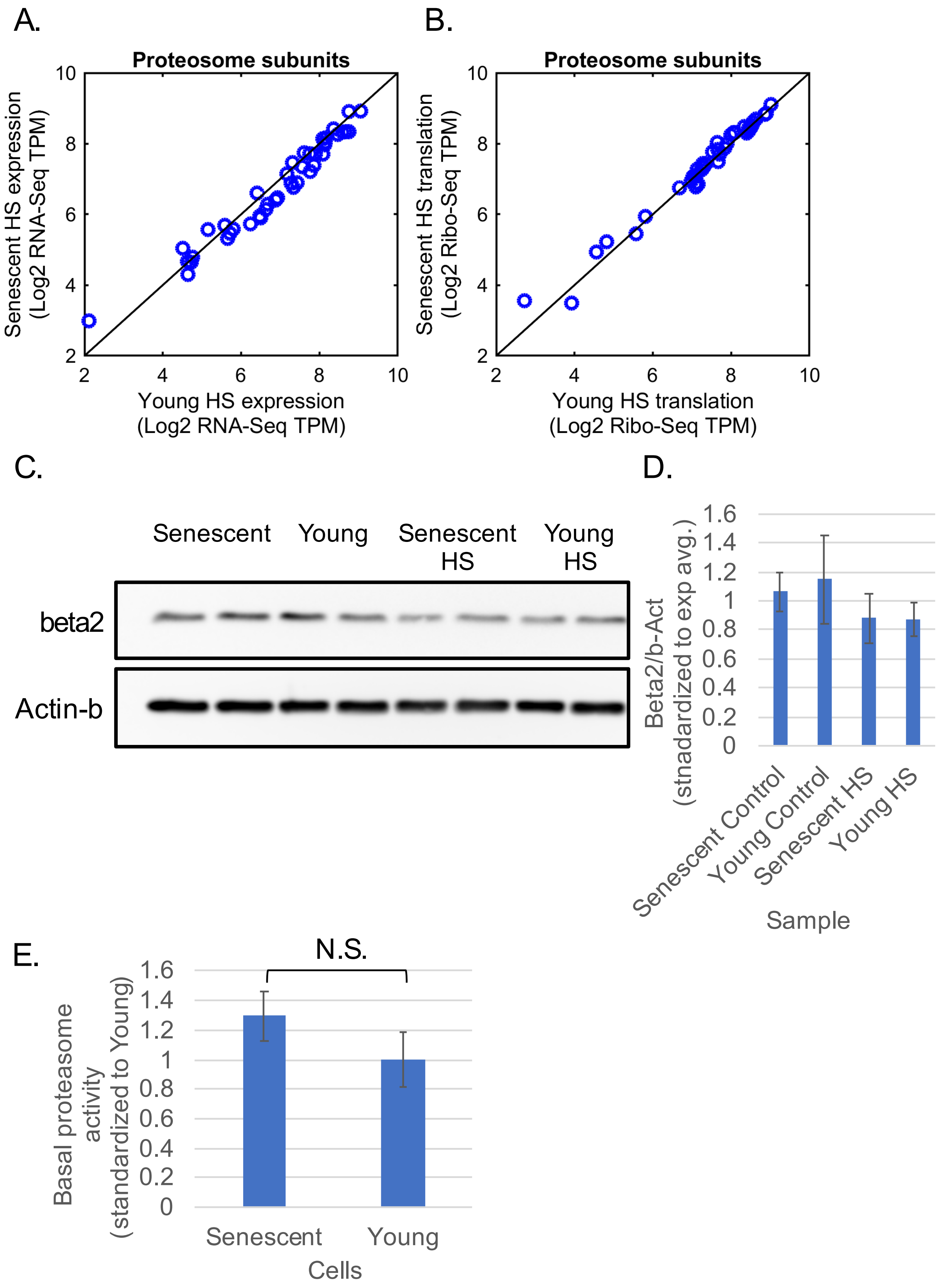
A.
Proteosome subunits
Senescent HS expression (Log2 RNA-Seq TPM)
Young HS expression (Log2 RNA-Seq TPM)
B.
Proteosome subunits
Senescent HS translation (Log2 Ribo-Seq TPM)
Young HS translation (Log2 Ribo-Seq TPM)
C.
Senescent
Young
Senescent HS
Young HS
beta2
Actin-b
D.
Beta2/b-Act (stnadardized to exp avg.)
Senescent Control
Young Control
Senescent HS
Young HS
Sample
E.
N.S.
Basal proteasome activity (standardized to Young)
Senescent
Young
Cells

**Figure S8: Deterioration of proteasome function in stressed senescent cells.**

(A-B) As in Fig. 7A,B, mRNA and translation levels of proteasome subunits are unchanged between young HS and senescent HS cells. (C-D) The protein levels of the beta2 subunit of the proteasome is unchanged between young and senescent cells. (D) quantification of beta2 levels (normalized to b-Actin) was performed using Fiji. Bars represent mean and std of 4, 5, 3 and 5 biological replicates for senescent, young, senescent HS and young HS cells respectively. None of the samples showed a significant change compared to the others (using t-test). (E) Proteasome activity assay (see Methods), bars represent mean and std of 7 biological replicates, data was normalized to the average of the young sample in each independent experiment in order to compare between different experiments. Basal levels showed no difference between young and senescent cells (using t-test, p=0.17).

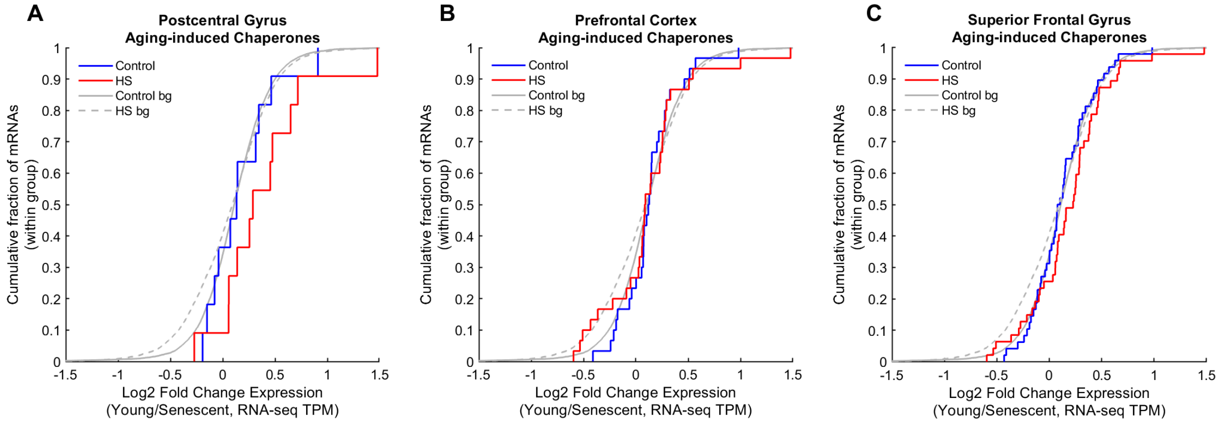


**Figure S9: Chaperones induced in aged brains do not show proteostasis decline behavior.** (A-C) Age-induced chaperones were taken from Brehme et al. (1). as defined for three different tissues. CDF plots depict the log2 fold changes expression of this group of genes in Young/Senescent cells. These chaperones were not significantly induced in heat shock in young cells more than in senescent cells, and therefore the HS curve is not significantly shifted compared to the background, or compared to the control curve (all p-values > 0.01). This is in contrast to aging-repressed chaperones defined in these three tissues by Brehme et al. (1), shown in Fig. 8.

**Table S1: Gene expression and translation data.**
Gene expression (RNAseq, denoted "rna") and translation (Riboseq, denoted "fp") values, as specified in the Methods, for all human genes. Additionally, the table specifies the p-value for differential expression in each sample (DESEQ2 output, denoted "DSeq_pval"), and if the gene was included as a differentially expressed by RNA expression or translation, a cluster number is denoted (otherwise cluster number is 0), in accordance with the cluster numbers in the supplementary figures.

**Table S2: mRNA expression and translation clusters.**
RNAseq (A) and translation (B) cluster numbers and colors (matching the colors in Fig. S6I), together with number of genes and the cluster behavior. Cluster numbers are in accordance with the cluster numbers in the supplementary figures.

**Table S3: Pathway enrichment analysis results for expression and translation clusters.**
Enriched pathways and annotations for each of the RNA expression (A) and translation (B) clusters. See Methods for pathway functional enrichment analysis.

**Table S4: Senescence-enhanced HS translational repression cluster - functional enrichment analysis.**

**Table S5: Manually curated chaperones list.**
Used in Fig. 2B,C, S3A,B.
